# Reading ability in both deaf and hearing adults is linked to neural representations of abstract phonology derived from visual speech

**DOI:** 10.1101/2025.11.09.684749

**Authors:** Samuel Evans, Cathy J. Price, Jörn Diedrichsen, Tae Twomey, Indie Beedie, Maggie Fraser, Mairéad MacSweeney

## Abstract

Reading is central to academic and vocational success. Some deaf children face reading challenges due to limited access to spoken or signed language. Robust phonological representations are key to reading development in hearing children. Spoken language phonology may be one of many contributors to reading development in deaf children. Indeed, speechreading ability correlates with reading skill in both deaf and hearing individuals, suggesting it is linked to reading development regardless of hearing status. Further support for this hypothesis would be provided by evidence that similar neural representations of speech phonology are evoked by visual speech and other language forms (auditory speech and text), and that these neural representations are related to reading proficiency. We used fMRI and Representational Similarity Analysis (RSA) to identify shared neural representations of spoken language phonological structure. A group of deaf adult participants (N=22), with a mixture of sign language and spoken language backgrounds and reading abilities, were presented with single lexical items as visual speech and dynamic text (cursive text, revealed letter-by-letter to promote a phonological reading strategy). Adult hearing participants (N=25) were presented with the same words, but as visual speech and auditory speech. Shared neural representations of phonological structure of English words were found in each group in the superior and middle temporal cortex (STC/MTC) and these abstract representations were more similar across different language forms in better readers. Our data provide neurobiological evidence of the contribution of visual speech to abstract phonological representations of spoken language, that relate to reading proficiency, in both deaf and hearing adults.

**Significance Statement:** Reading is an essential skill, yet some deaf children face reading challenges due to reduced access to signed or spoken language. In hearing children, successful reading depends on abstract phonological representations, but whether spoken language phonology relates to reading in deaf individuals remains unclear. Using fMRI and RSA, we show that deaf and hearing adults recruit neural representations of phonology that are shared by visual speech and other language forms (visual/auditory speech in hearing; visual speech/dynamic text in deaf) in the superior and middle temporal cortex. Critically, greater cross-modal alignment of neural representations of phonological structure was associated with better reading in both groups. These findings provide neurobiological evidence that visual speech contributes to phonological representations that relate to reading, regardless of hearing status.

## Introduction

The ability to read is essential for acquiring knowledge and to participate fully in educational, professional, and social life in industrial societies. Some deaf children face challenges in learning to read, as a result of limited access to language - spoken or signed - during early development (1–4). Most hearing children learn to read by linking speech sounds to letter forms. A key factor in this process is the ability to access and manipulate well-specified, abstract ‘phonological representations’ of spoken language. This skill is an important predictor of reading in hearing children (5) and interventions designed to strengthen phonological awareness are effective in remediating reading difficulties (6).

In deaf children, numerous studies show evidence of positive correlations between phonological awareness of speech and reading proficiency (7–9), although this association has not always been observed (10). Indeed, the importance of phonological awareness in reading development in deaf children is hotly debated (2, 11) and evidence suggests that there are multiple pathways to reading success for deaf readers (12–14). For those deaf people for whom awareness of the phonological structure of speech is associated with reading proficiency - how might they acquire this? One way is through visual speech (speechreading/ lipreading), which is the primary means by which spoken language is accessed by all severe-profoundly deaf people, whether signers or non-signers. Positive associations between speechreading and text reading have been observed in deaf children and adults, and hearing children (15–19). It is likely that this relationship operates at many linguistic levels – not only lexical but also sublexical. That is, visual speech provides a visual source of information about spoken articulations that helps to establish better specified phonological representations and/or promotes greater awareness of phonological structure. In hearing individuals, visual speech likely plays an important but more supplementary role, providing redundant and complementary information that supports auditory perception (20). Indeed, behavioural studies in deaf and hearing children suggest that the observed relationship between visual speech and reading ability is related to phonological awareness, such that visual speech may contribute to phonological representations, which may in turn support learning to read (15, 18, 21). The current study used fMRI and Representational Similarity Analysis (RSA) to seek neurobiological support for this proposal by examining whether visual speech evokes neural representations of phonological structure that are shared between visual speech and other language forms – specifically auditory speech in hearing adults and dynamic text (i.e., cursive text, revealed letter-by-letter) in deaf adults. We also assessed whether these representations relate to reading ability. We used dynamic, rather than static, text to promote sublexical reading (22).

The superior and middle temporal cortex (STC/MTC) is engaged by phonological processing across a range of language forms, including auditory speech perception and production (23–25), speechreading (26, 27), text reading (28–30) and audio-visual integration (31–33). These regions are also responsive during speechreading in people who are severely-profoundly deaf (34). This broad responsiveness to different language forms suggests that the STC/MTC may support ‘modality independent’ neural encoding of speech phonology. An alternative view is that different language forms evoke overlapping but distinct patterns of activity suggesting parallel but modality-specific processing.

Strong evidence for modality *independent* representations has been provided by showing that neural patterns evoked by different language forms share common representational geometry (35). This approach has been used to demonstrate similar encoding of semantic information across the visual and auditory modality for auditory speech and sign language in hearing signers (36) and between auditory speech and text in hearing spoken language users (37). It has also been used to test for shared phonological representations of auditory and visual speech in spoken language users, although findings in this area are equivocal. For example, Keitel et al., (2020) found evidence for co-located but modality specific representations of heard and seen words in the MTC, and other regions. By contrast, Van Audenhaege et al., (2025) found that neural patterns for the same auditory and visual speech syllables were shared in bilateral posterior superior temporal sulcus and left somatomotor cortex. Neither of these studies addressed whether common representations of phonology were related to text reading ability – a key academic and cognitive skill. Furthermore, neither study involved deaf participants for whom the role of phonology of speech in reading is unclear.

In summary, we predicted that representations of phonological structure shared across language input forms would be found in the bilateral STC/MTC in both hearing and deaf groups. We also predicted that better text readers would have neural representations of phonology that were more similar across language forms. Such evidence would provide neurobiological support for the hypothesis that visual speech contributes to abstract representations of phonology that relate to reading in both deaf and hearing people.

## Results

A group of participants with typical hearing, preselected for their good speechreading ability, were scanned with functional Magnetic Resonance Imaging (fMRI) while attending to the same eight words (see Fig. 1A, left) produced by two different speakers in both visual speech and auditory speech conditions. A group of severely-profoundly deaf participants were also scanned while attending to the same words presented as visual speech or dynamic text (i.e., cursive text, revealed letter-by-letter) in two different fonts (see Fig 1A, right).

**Fig 1.**
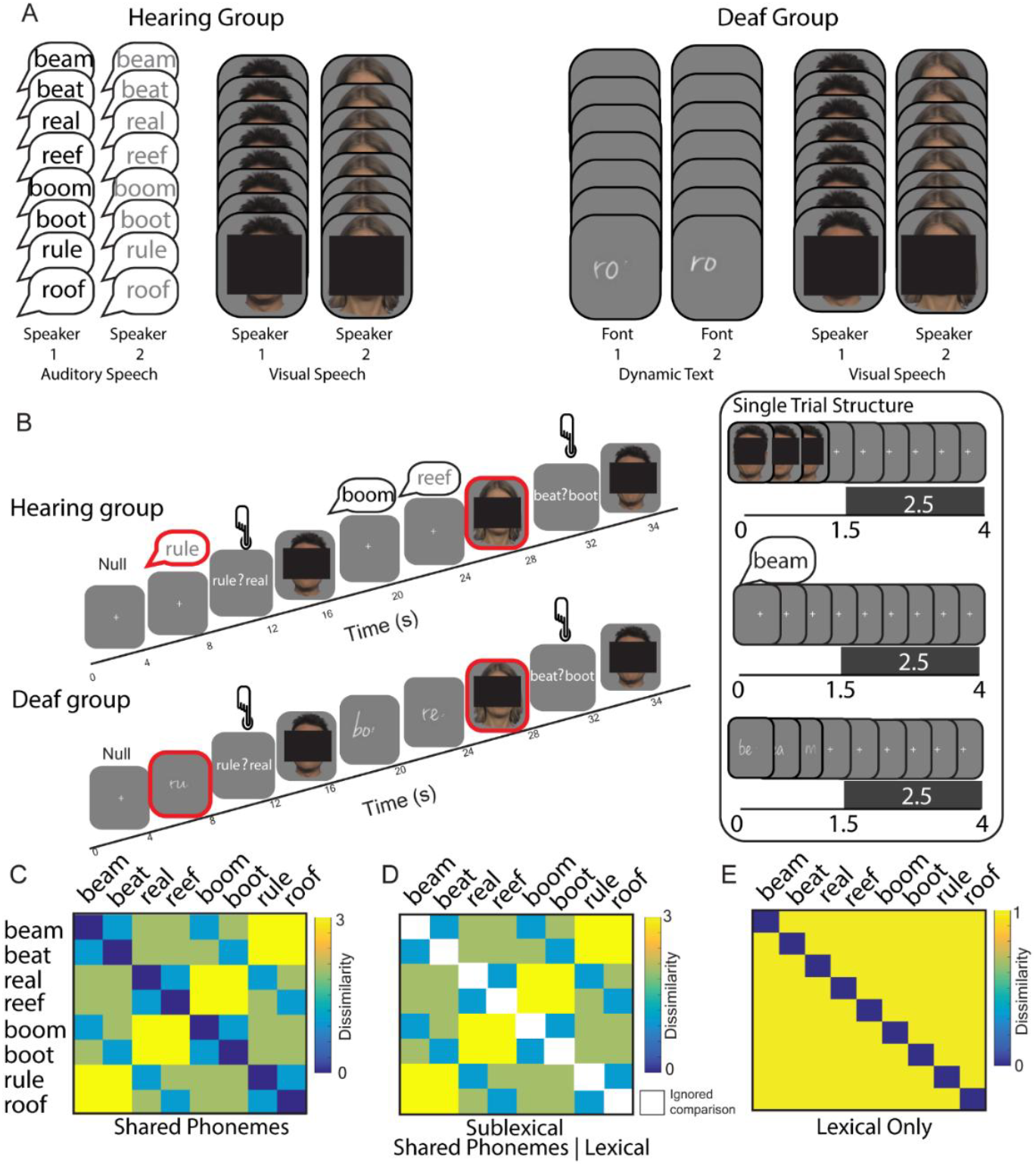
fMRI Scanning Paradigm and RSA Models. (A) Hearing and deaf participants were presented with the same eight words. These were presented as visual speech and auditory speech for hearing participants and visual speech and dynamic text (cursive text, revealed letter by letter) for deaf participants. (B) Participants engaged in an occasional one-back forced choice word identification task during scanning in which they had to select the word they had just seen from two alternatives. A target trial, highlighted with a red outline on the figure only, preceded decision trials in which a button press was required. Using a fast-sparse scanning sequence, all stimuli were presented in a 1.5 second silent gap between 2.5 second volume acquisitions so that the phonetic content of the auditory speech could be heard clearly by hearing participants. (C) Shared Phonemes model - quantified the dissimilarity between words based on the number of shared phonemes, without distinguishing lexical and sublexical dissimilarity. (D) Sublexical model (Shared Phonemes | Lexical) - quantified dissimilarity based on the number of shared phonemes but excluded dissimilarity comparisons of each word with itself. (E) Lexical model - quantified dissimilarity on the basis of whether the word was the same or not. This model predicts that each word is most similar to itself and maximally and equally dissimilar to all other words. Note that faces have been obscured in section A due to BioRxiv policy – the videos included facial features in the experiment.

When measured outside the scanner, text reading scores were lower in the deaf compared to hearing group, but the reverse was true for speechreading, which was better in the deaf group (see Participants section & SI Appendix, Fig S1). There was a significant positive relationship between speechreading and text reading in the deaf group, but not the hearing group. During scanning, both groups were highly accurate on an occasional 2-Alternative Forced Choice (2AFC) task in which they had to identify the last seen or heard word (> 90% accurate, SI Appendix, SI Methods).

### Neural representations of abstract speech phonology

Prior to the main analysis, we tested for evidence of abstract neural representations of sublexical structure *within each individual stimulus* by testing for shared representations across speakers and/or fonts (SI Appendix, SI Results). This provided evidence that sublexical structure was encoded in the STC/MTC neural patterns for each stimulus type in a manner that was abstracted from lower-level sensory features. We then conducted our key analyses, testing for shared representations of sublexical structure *across stimulus types, within each group*.

#### Hearing participants

Regions that encode shared representations of phonology *across-stimulus* type also need to encode phonology *within* both stimulus types. Using Representational Similarity Analysis (RSA), we estimated the activity patterns elicited by each trial and calculated the distances between activity patterns within searchlight regions across the whole brain (see method). We then identified clusters with reliable non-zero within-stimulus representational distances (i.e., both within visual speech *and* within auditory speech, see Fig 2B, grey boxes). This identified six clusters, in [1] the left STC/MTC and [2] right STC/MTC, [3] right V1-V3 and [4] left inferior frontal gyrus, [5] right posterior middle temporal gyrus and [6] left V1-V3 (Fig 2A, see also SI Appendix Table S4).

**Fig 2.**
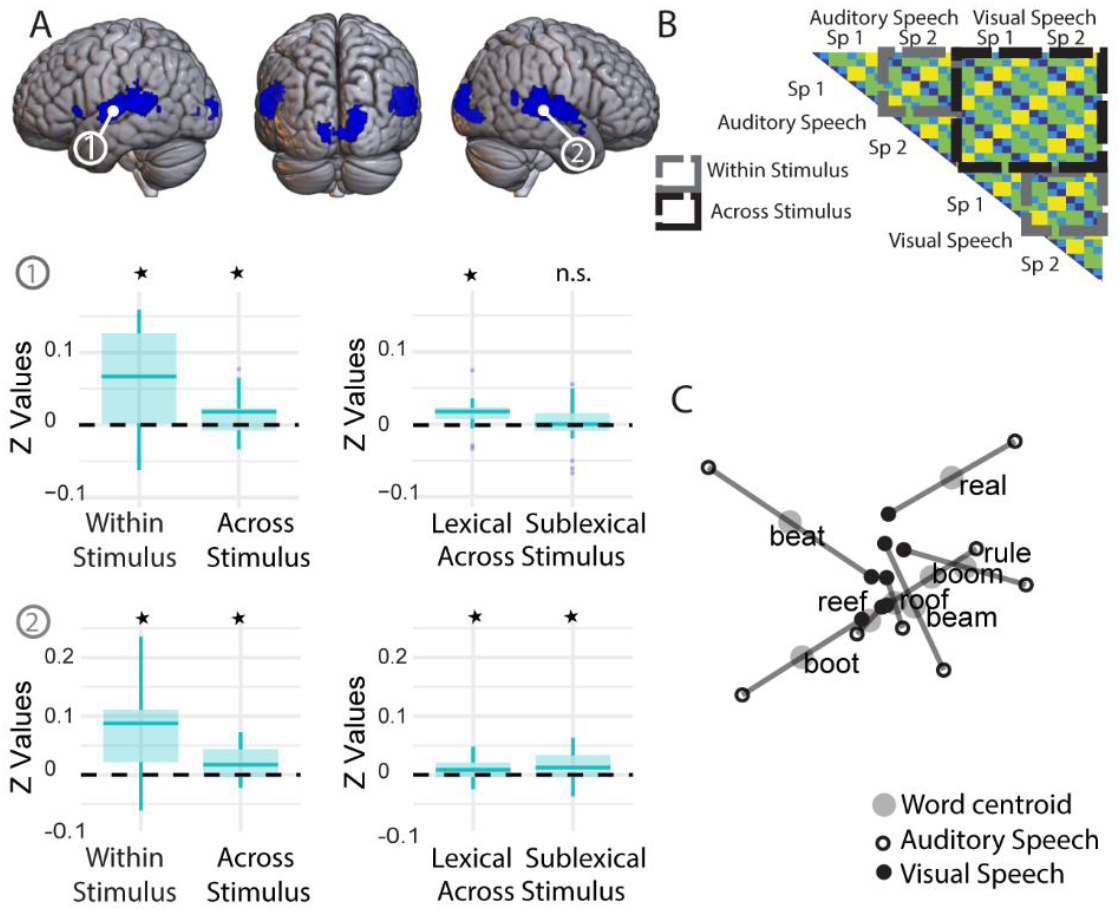
Hearing participants - Searchlight clusters with non-zero within stimulus type distances in hearing participants, rendered on the MNI brain and thresholded at p < 0.001 peak uncorrected, q < 0.05 FDR cluster corrected. (A) Neural patterns in bilateral STC/MTC were a significant fit to the Shared Phonemes model in the within and across-stimulus distances (for auditory and visual speech). Only the right STC/MTC – cluster 2 – was a significant fit to the Sublexical model within and across-stimulus type. *Indicates statistical significance correcting for multiple comparisons. (B) In regions with positive distances within-stimulus, we tested for a fit to the models in the within-stimulus (grey boxes) and across-stimulus distances (black box). (C) Multi-Dimensional Scaling (MDS) visualisation of the neural patterns for the hearing group in the right STC/MTC showing the similarity between the neural patterns for each word, in each stimulus type, in a 2D representational space.

The clusters in the bilateral STC/MTC were mainly in the STC, but also extended posteriorly into the middle temporal gyrus. There was only a significant fit to the Shared Phonemes model (Fig 1C) in both the within and across-stimulus distances in these regions (Fig 2B). The fit to this model could be driven by lexical-semantic and/or sublexical-phonological processes. To ensure that the model fit was driven by shared representations of sublexical-phonological structure we further assessed the fit to a *Sublexical model* (Shared Phonemes model | Lexical, Fig 1D) which quantified dissimilarity between words on the basis of the number of shared phonemes, but excluded distances between words of the same identity. This model was a significant fit in the right STC/MTC [peak at 57 −31 5] in both the within and across-stimulus distances (Table 1, extended table in SI Appendix S5) and within-stimulus when the auditory and visual speech were tested separately (both ps <= 0.029). This provided clear evidence of shared sublexical representations of auditory and visual speech in the right STC/MTC (see Fig 2C for MDS plot).

**Table 1.**
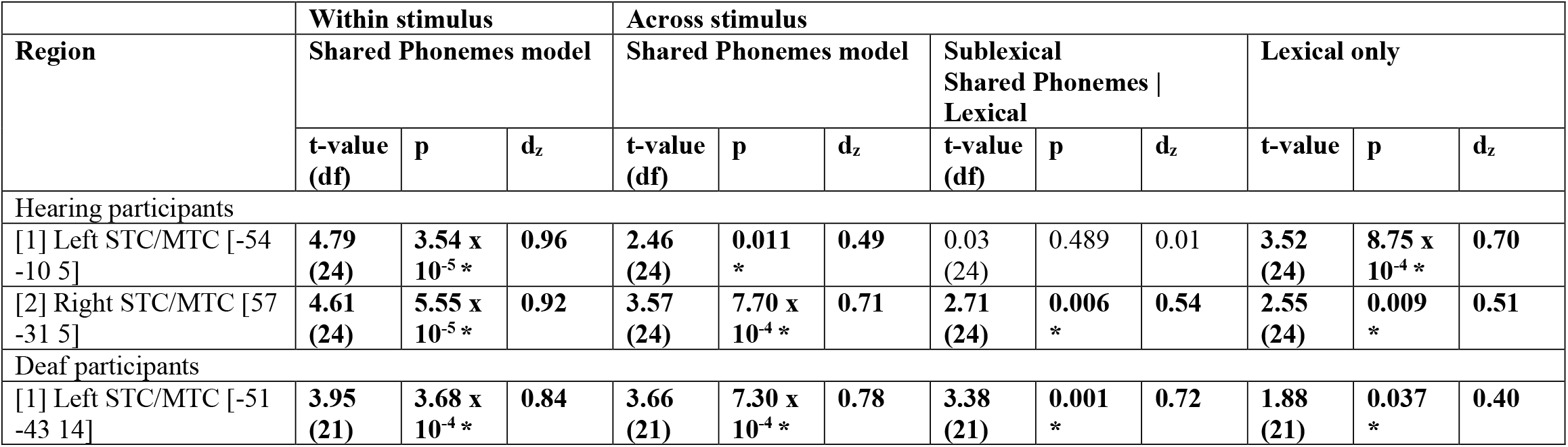
Model fits in the STC/MTC to the within and across stimulus distances in hearing and deaf participants. Tests for fits to the Shared Phonemes model were only conducted in the across-stimulus distances, if there was a fit to the model in the within-stimulus distances. The Sublexical and Lexical models were only tested in the across-stimulus distances if there was a fit to the Shared Phonemes model in the across-stimulus distances. Multiple comparisons were Bonferroni corrected for the number of clusters in which each model was tested. **\*Indicates statistically significant fit corrected for multiple comparisons**. See SI Appendix, Table S5 for model fits in all clusters, including those that were not a significant fit to the Shared Phonemes model within stimulus type, in areas outside the MTC/STC.

In the left STC/MTC [peak at −54 −10 5], there was not a strong fit to the Sublexical model, but there was a fit to a *Lexical model* that predicted that words of the same identity were least dissimilar to one another and maximally and equally dissimilar to the other words in the set (Fig 1E). There was no evidence of a difference in strength of fit between the Lexical and Shared Phonemes models (t (24) = 0.03, p = 0.977, d_z_ = 0.01). This was indicative of abstract lexical-semantic rather than sublexical-phonological representations.

Given that we had also expected a fit to the sublexical model in the left STC/MTC in hearing participants (see Van Audenhaege et al., 2025), we ran an additional searchlight analysis directly testing for a fit to the sublexical model within this cluster. A number of small clusters of voxels were identified (SI Appendix Fig S8) as a fit to the Sublexical model but only when lowering to an uncorrected peak level threshold of p<0.005.

#### Deaf participants

We adopted the same analytical approach for analysing the data from the deaf group. A searchlight identified eight clusters with reliable non-zero within-stimulus distances for dynamic text and visual speech (i.e., both within visual speech *and* within dynamic text, Fig 3B). These clusters were found in [1] left STC/MTC, [2] bilateral V1-V3, extending into left middle temporal cortex, [3] right STC, [4] right MTC, [5] right inferior frontal gyrus, [6] left precuneus, [7) left inferior frontal gyrus and [8] right middle occipital gyrus (Fig 3A, see also SI Appendix, Table S4). The left STC/MTC cluster was centred in the superior temporal sulcus extending into the superior and middle temporal gyrus [peak at −51 −43 14]. This was the only region to show a significant fit to the Shared Phonemes model in the within and across-stimulus distances (Table 1, with extended table in SI Appendix Table S5). There was also a fit to the Sublexical model within and across stimulus, and when the dynamic text and visual speech distances were tested separately (both ps <= 0.019). This provided clear evidence for shared sublexical representations of visual speech and dynamic text in the left STC/MTC (see Fig 3C for MDS plot).

**Fig 3.**
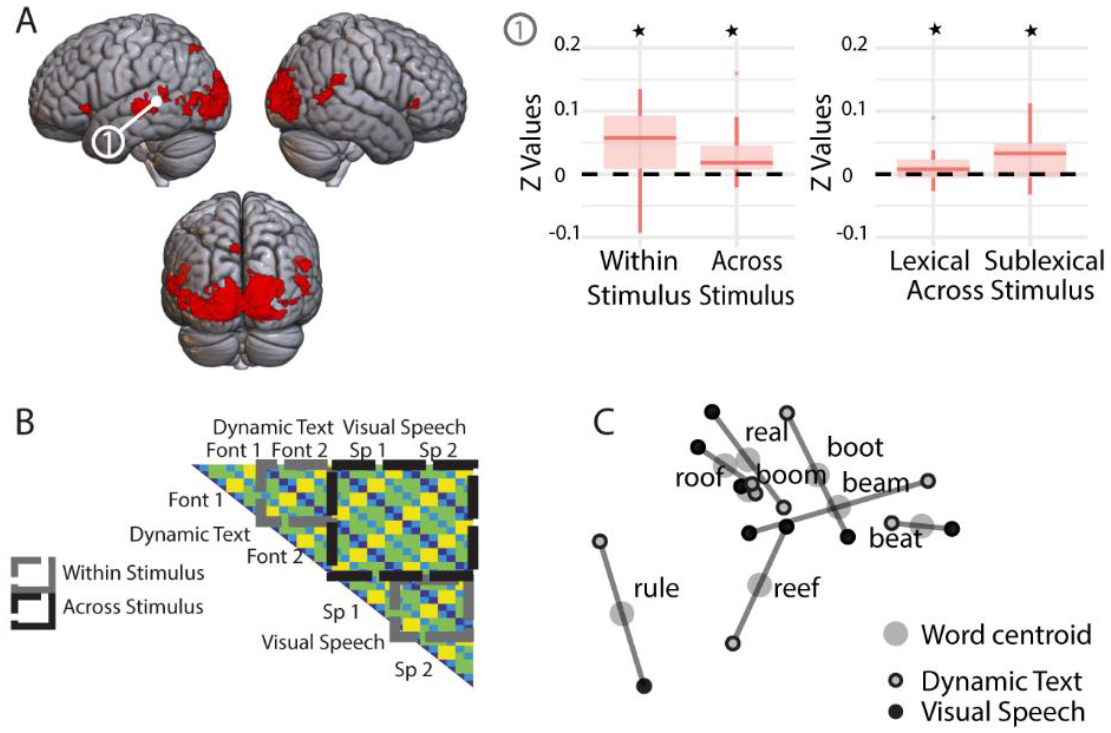
Deaf Participants - Searchlight clusters with non-zero within stimulus type distances in deaf participants, rendered on the MNI brain and thresholded at p < 0.001 peak uncorrected, q < 0.05 FDR cluster corrected. (A) Only neural patterns in the left STC/MTC – cluster 1 - were a fit to the Sublexical model in the within and across-stimulus distances (for dynamic text reading and speechreading). *Indicates statistical significance correcting for multiple comparisons. (B) In regions with reliable within-stimulus distances, we tested for a fit to the models within (grey boxes) and across-stimulus type (black boxes). (C) Multi-Dimensional Scaling (MDS) visualisation of the neural patterns for the deaf group in the Left STC/MTC showing the similarity between patterns for each word, in each stimulus type, in a 2D representational space. Note that the MDS solutions were computed independently for each group. Therefore, only *relative* distances within each MDS plot are interpretable.

We further tested whether there were differences in the strength of the encoding of abstract phonemes in the different segment positions of the words (initial consonant, vowel, final consonant) in the regions shown to encode abstract sublexical representations in each group (SI Appendix, Fig S9). There was no evidence of a difference in strength of encoding across phoneme position (hearing: F (2,48) = 0.720, p = .492; deaf: F (2,42) = 1.613, p = .211), suggesting that all phonemes were represented with equivalent strength in both groups in these regions.

### Relationship to reading

To test for the functional relevance of these neural representations of phonological structure that were shared across stimulus form, we correlated the fit to the Shared Phonemes model in the across-stimulus distances with reading scores collected outside the scanner. We tested this in the regions shown to encode sublexical representations across stimulus type in each group separately. In hearing participants in the right STC/MTC cluster, there was a positive relationship between the fit to the Shared Phonemes model across stimulus type (for visual and auditory speech) and reading scores (r (24) = 0.434, p = 0.015, one-tailed test, Fig 4A). In deaf participants in the left STC/MTC cluster, there was a significant correlation between the model fit to the shared phonemes model across stimulus type (for visual speech and dynamic text) and reading scores (r (21) = 0.463, p = 0.015, one-tailed test, Fig 4B).

**Fig 4.**
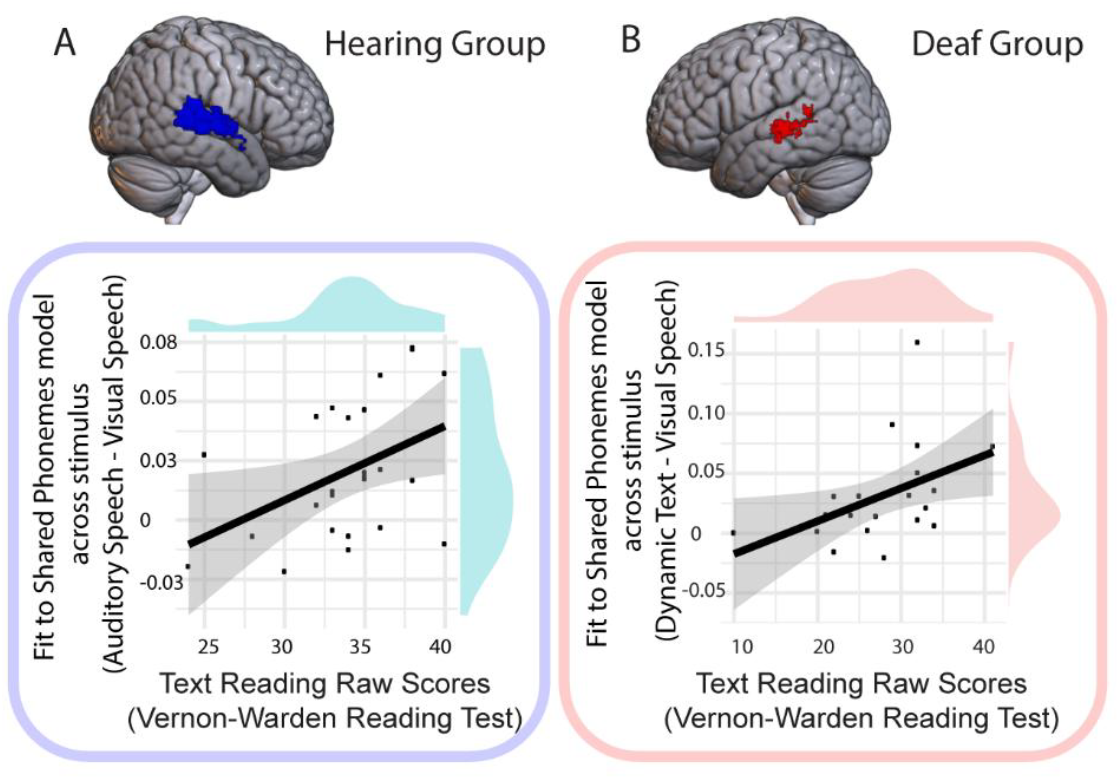
Relationship between abstract speech phonology and reading proficiency. (A) In hearing participants in the right STC/MTC, there was a positive relationship between the strength of encoding of abstract neural representations of phonological structure (i.e. shared between auditory and visual speech) and text reading measured outside the scanner. (B) In deaf participants in the left STC/MTC, there was a positive relationship between the strength of encoding of abstract neural representations of phonological structure (i.e. shared for dynamic text and visual speech) and text reading measured outside the scanner.

To summarise, we found neurobiological evidence of abstract representations of speech phonology in the left and right STC/MTC respectively in deaf and hearing individuals. These shared neural representations were functionally related to reading proficiency in both groups, such that better readers had neural patterns in which the representational geometry across-stimulus type was more closely aligned with phonological structure.

## Discussion

Behavioural associations between speechreading and text reading ability in deaf and hearing children suggest that speechreading may contribute to reading development across populations (15–17). The extent to which spoken language phonology contributes to reading in deaf people remains unclear (2, 11). Understanding how neural representations of visual speech relate to neural representations of text and to reading skill can provide a new perspective on this issue. We provide neurobiological support for a relationship between abstract phonological representations of speech and reading proficiency in deaf and hearing adults by identifying evidence of neural representations of phonology shared across language input forms in the STC/MTC in each group. Furthermore, we show that individuals were better text readers if their neural representations of spoken language phonology were more similar between the language forms that they were exposed to.

### Shared neural representations of phonological structure for auditory and visual speech and their relation to reading in hearing adults

We found evidence for shared neural representations of phonology for auditory and visual speech in hearing adults in the STC/MTC. This stands in contrast to Keitel et al. (2020) who found that word identity could be decoded from auditory and visual speech separately in the bilateral middle and inferior temporal cortex, but not *across* auditory and visual speech, indicative of co-located but distinct representations. The difference in findings with our study likely reflects design choices. Keitel et al. (2020) reduced the intelligibility of auditory speech by adding background noise to equate intelligibility with the visual speech condition. Nevertheless, there was still much greater variation in accuracy in the visual speech, than auditory speech, condition. Lower intelligibility across both conditions and the greater variability in visual speech accuracy (ranging from ~30% to ~95% accuracy across participants) may have reduced the likelihood of finding robust shared representations. By contrast, we maximised the intelligibility of our stimuli by using a small set of predictable, highly speechreadable words and pre-selecting hearing participants who were good speechreaders, ensuring the best chance of finding shared representations across auditory and visual speech. Whilst this was a successful strategy, future studies involving more naturalistic communicative interactions, including a wider set of words, is necessary to assess the extent to which our findings generalise to other communicative situations and participant groups.

Our results are more closely aligned with the study of Van Audenhaege et al., (2025) who found evidence for shared representations of non-word syllables for auditory and visual speech in the posterior bilateral STC. Using an RSA model-based approach, we were able to extend these findings by differentiating between lexical-semantic and sublexical-phonological representations, and by relating shared sublexical representations to reading performance. We only found strong evidence for shared sublexical-phonological representations in the right STC/MTC and did not find equivalent evidence for shared sublexical representations in the left STC/MTC. The finding of lexical representations in the left STC/MTC is consistent with studies showing a left dominant response to meaningful speech (40). However, a model-based searchlight analysis identified some sensitivity to sublexical-phonological structure in subregions of the left STC/MTC, at an uncorrected level. Hence, it may be that the absence of strong evidence of shared sublexical representation in the left STC/MTC reflected our methodological approach, rather than a more fundamental difference in function. Future studies are necessary to explore this further.

Our novel finding, that shared neural representations of phonological structure evoked by visual and auditory speech are linked to reading ability in hearing individuals extends, previous work. Phonological awareness is a strong predictor of reading success in hearing children (6).

These abilities rely on accessing high quality phonological representations of spoken language stored in memory. The role of visual information in establishing these representations has often been overlooked in preference of an audiocentric perspective in phonology research (41).

Nevertheless, visual speech plays an integral role in speech perception. Speech is usually both seen and heard, and when visual speech is available, it is automatically and obligatorily integrated (42). Visual speech cues can facilitate auditory perception by providing access to redundant and complementary information that supports speech understanding, particularly when listening is difficult (20). Indeed, sensitivity to visual speech develops early, with infants showing awareness of the congruence between lip movements and speech sounds soon after birth (43) and neural specialisation for visual speech and audiovisual integration developing over the first few months of life (44–46). Our data provide neurobiological evidence of a link between abstract neural representations of phonology, shared between auditory and visual speech, and reading ability, consistent with a role for visual speech in establishing multimodal phonological representations (47). This provides a neurobiological basis for the positive association between speechreading and text reading in hearing children (15, 18). It is also consistent with a recent study in hearing children showing that univariate neural activity in the STC is associated with both visual speech perception and rhyme judgments of written words (48). Studies showing that speechreading skills are “trainable” in hearing children (49) and that children with reading and language disorders benefit to the same degree from audio-visual speech in noisy conditions as their typically developing peers (50, 51), suggest that reading instruction with a focus on speechreading may show promise in supporting struggling hearing readers. Intervention studies are necessary in hearing children to assess the efficacy of this approach.

### Shared neural representations of phonological structure for dynamic text and visual speech and their relation to reading in deaf adults

Given deaf children’s reduced access to sound and the importance of phonological awareness to reading success in hearing children, this has led to the assumption that improving speech-based phonological skills in deaf readers should be the focus of reading interventions (11). Indeed, some profoundly deaf children show above chance judgments of rhyme and syllable structure (52) and some young deaf adults show patterns of reading performance that suggest the implicit use of speech phonological codes (53), indicative of a degree of phonological awareness. However, it has also been argued - by ourselves and others - that phonological skills are not necessarily a requirement for skilled reading in deaf readers, some of whom can achieve high levels of literacy via non speech-phonological routes (12–14, 54). This is a perspective consistent with connectionist models of reading that emphasise the importance of different forms of representation in reading (55) such that deaf readers may rely greatly on connections between orthographic and semantic processing, in addition to drawing on other forms of representation derived from sign language (56). However, we note that these different perspectives on the role of spoken language phonology in deaf readers are not necessarily incompatible. Evidence suggests that the extent to which deaf readers use speech phonology during reading, versus other linguistic sources, likely depends on multiple factors, including language and sensory experience (12) and task demands (54).

fMRI studies of phonological processing in deaf early signers suggest that they may rely more on non-phonological, semantic-orthographic routes to reading (57–61). When asked to make explicit phonological judgments about spoken language (e.g., judging number of syllables or whether the English labels for the pictures ‘chair’ and ‘bear’ rhyme?), both deaf and hearing participants activate inferior parietal and temporoparietal cortex (57, 62, 63). However, deaf readers often activate these regions more and are less accurate on these tasks than hearing readers (57, 62) or exhibit coarsely tuned neural representations of speech phonology measured with repetition-suppression (59). By contrast, neural responses in the left ventral occipital cortex (sometimes referred to as the visual word form area (64)) display fine-tuned orthographic selectivity in deaf readers (59, 60). Together, these findings have been used to argue that deaf readers rely more on precise orthographic representations than on their comparatively coarser (speech) phonological ones (61).

Rather than using explicit phonological judgments (which draw on metacognitive skills) or using phonological manipulations within a single modality using repetition-suppression, we identified regions in which neural representations of phonology were shared between visual speech and dynamic text. We used a more implicit measure of phonology, asking participants on occasional trials (~13% of trials) to select the previously seen real word from a two-alternative forced choice. This required minimal memory and task demands, whilst allowing us to monitor that participants were attending to the stimuli. Evidence of phonological processing was inferred from common activity patterns evoked by both dynamic text and visual speech that reflected the spoken language phonological structure of our stimuli. Our finding of shared neural representations for visual speech and dynamic text in the left posterior STC/MTC is consistent with previous univariate studies of reading and visual speech processing in deaf and hearing individuals that associate activity to this region (28, 30, 32, 65–67). Indeed, the left posterior STC has been shown to be more strongly activated during reading in more proficient, as compared to less proficient, deaf readers suggesting an association between activity in the left STC and reading success (66). However, the region that we identified is more anterior and inferior to the temporo-parietal and inferior parietal regions identified in the studies of reading involving explicit phonological judgements reviewed above. This may reflect our approach of looking for regions of shared representations across language forms which may be more closely associated with speech processing in the STC.

How might the use of dynamic text have influenced our findings? When reading dynamic text, participants are unable to see the whole word during stimulus presentation, as letters appear and disappear over time. This prevents whole word recognition strategies and encourages assembly of segmental phonology. In deaf participants, we found shared neural patterns for dynamic text and visual speech that were aligned with the segmental phonological structure of each word in a similar region of the left posterior STC/MTC that was activated by dynamic text in hearing individuals in our previous study (22). This, we argue, is consistent with the interpretation that deaf participants were accessing speech phonological structure when presented with these stimuli. However, it does not follow, and we do not argue, that deaf people always draw on speech phonology when reading. Indeed, we have shown, the extent to which deaf individuals make use of speech phonology when reading likely differs greatly within and between individuals, depending on intrinsic and extrinsic factors (54). Deaf participants rely on multiple reading strategies that include drawing heavily on orthographic pattern recognition, morphological knowledge, semantic and contextual understanding, and, for signers, sign-based linguistic representations, including fingerspelling (61, 68). In this study, we deliberately encouraged use of speech phonology by using dynamic text, in addition to visual speech.

Dynamic text is an unusual format for text that is not encountered outside of the laboratory. Further studies using static text are necessary to understand the extent to which our findings generalize to other reading contexts.

It seems unlikely however that the data from deaf participants in the current study can be explained by non-phonological reading strategies. An explanation based on shared orthographic processing is unlikely, given that letter representations are not available from watching the face during the visual speech condition. Moreover, an orthographic recoding account would predict shared activity in the ventral occipitotemporal cortex (vOTC), which we did not observe. Equally recoding to a sign-based representation would predict a fit to the lexical and not the sublexical model. A further alternative could be recoding to a fingerspelling-based representation, however, again, if this was the case, shared patterns would be expected in the vOTC (67) rather than STC/MTC. Given these arguments, the most parsimonious explanation is that our findings reflect abstract representations of speech phonology in deaf readers that are similar for visual speech and dynamic text.

It is not clear how the language experiences of the deaf participants in the current study influenced our findings. Previous fMRI studies with deaf readers have mainly tested exclusively deaf native and early signers (57, 59, 60). Native signers form only 5-10% of the deaf population (69). In the current study, we intentionally recruited a very diverse group of severely-profoundly deaf adults, representative of the wider deaf population. The majority of the deaf participants in the current study were born to hearing parents, hearing aid users and learned BSL after the age of five years, which they used as their preferred language in adulthood. Although many deaf individuals had a reading level equivalent to that of the hearing participants, on average the deaf readers as a group in the current study did not score as highly on the reading task as the hearing group. Future studies with sufficient sample sizes to compare deaf early and late signers, and low and high skilled readers, are necessary to understand how these factors influence associations between abstract neural representations of speech phonology and reading ability.

Our findings show that visual speech contributes to neural representations of abstract speech phonology that are linked to reading ability in a relatively heterogeneous group of deaf individuals. A recent RCT trial provided evidence that visual speech training influences phonological representations in deaf children (21). However, this study did not demonstrate that this training transferred to reading improvements, although the period of training was relatively short (8 hours over 3 months). Phonological based interventions should not be considered the ‘sine qua non’ or ‘default’ for deaf children. Indeed, our findings are not supportive of an oral only approach to deaf reading education and we argue against a one size fits all policy that encourages phonological instruction over other reading strategies for deaf readers (68). Ongoing research, in our and other research laboratories, provide abundant evidence that early access to sign language can play an important role in scaffolding acquisition of spoken and written English (70, 71) and that sign language specific representations, such as fingerspelling, can facilitate acquisition of literacy skills (72, 73). Hence, a translanguaging approach tailored to each individual that makes full use of all the linguistic resources available to deaf readers is likely to be the most effective approach in strengthening links between speech phonology, semantic, orthographic, fingerspelling and sign-based representations, to support reading development (74).

### Integrating findings across deaf and hearing groups

Our data provides evidence for abstract neural representations of phonology to which visual speech contributes in both groups. The strength of these abstract phonological responses in STC/MTC are linked to reading in both deaf and hearing participants. We are mindful, however, of the difficulty of inferring the direction of causality in the relationships identified. It may be that speechreading abilities help to refine phonological representations that in turn support reading, that reading experience improves phonological awareness and speechreading abilities, or that they are reciprocally related (15, 18). It may also be the case that they are driven in part by shared latent variables related to broader linguistic abilities, such as syntactic, morphological or vocabulary knowledge, in sign or speech. For example, speechreading and English vocabulary contribute both shared and unique variance in predicting reading accuracy in hearing children (75). Further longitudinal brain imaging studies are necessary to unravel these relationships and associated neural mechanisms.

Our findings do not show that these abstract representations are necessarily the same in the two groups. Phonological processing is not a single unitary construct (76–78) and the exact phonological demands are likely to differ for the different stimuli presented to the deaf and hearing participants. Presenting different stimuli to deaf and hearing groups allowed us to test for direct evidence of speech-derived abstract phonological representations, using auditory speech, in the hearing group. This was not possible with deaf participants. Instead, we used dynamic text to encourage indirect access to speech-based phonology. The structure of abstract phonological representations of speech may also differ between groups for other reasons, given that they are built primarily from coarse grained visual speech inputs in deaf individuals but shaped by both auditory and visual speech inputs in hearing individuals. Additionally, cortical reorganization (79, 80) might further differentiate these neural representations.

It is notable that whilst the left inferior frontal cortex contained reliable within-stimulus representational structure in both deaf and hearing participants, this structure was not related to speech phonology as operationalized by the Shared Phoneme Model (and related models).

Similarly, we did not find phonological, or in fact any reliable representational structure, in the ventral occipital temporal cortex. This stands in contrast to recent evidence of co-located but modality specific representations of auditory non-words and visual speech (39) and auditory speech and text in this region (81). One explanation for these absences may be due to the use of real words and lack of explicit and challenging phonological task (63, 82). Future studies are necessary to adapt stimulus properties, task demands and representational models of phonology capable of identifying other representations used during language processing in these regions.

### Summary

We found evidence for neural representations of abstract speech phonology by assessing the similarity of neural representations evoked by visual speech, and other language forms, in deaf and hearing adults. These neural representations were related to reading outcomes in both groups. Our data provides neurobiological evidence for a relationship between abstract phonological representations of speech and reading proficiency in deaf and hearing adults. These data highlight the potential utility of visual speech in supporting struggling deaf and hearing readers. However, the direction of this relationship is still to be established and these findings should not be interpreted as supportive of an oral only approach to literacy education in deaf children. Rather they highlight the potential of visual speech to support reading in deaf readers, to an extent that is likely to vary based on a range of audiological and language background factors.

## Materials and Methods

### Participants

Ethical approval was granted by the UCL Research Ethics Committee and informed consent was obtained from all participants. Hearing participants were pre-screened to ensure a high level of speechreading ability (see SI Appendix, SI methods). Twenty-five right-handed typically hearing adults without known hearing, language or neurological impairments were scanned (N=25, mean age = 34 years; SD = 13 years, age range = 19-63; female = 13, male = 12).

The data from twenty-two severe-to-profoundly deaf right-handed adults were analysed (N = 22, mean age = 38; SD = 11 years, age range = 19-64; mean dBHL in better ear = 97; SD = 13, range in dBHL = 70-115; female = 10, male = 12, see SI Appendix, SI Methods and Table S1). All participants were deaf before the age of three years old. Ten participants learned BSL under 5 yrs old, 11 after 5 yrs old and 1 did not know. There was a range of self-rated BSL proficiencies (mean proficiency (scale 1-7) = 5.8, SD = 1.4, range = 2-7). The majority used BSL as their preferred language (Preferred language: BSL = 12; Sign Supported English = 6; spoken English = 4). Sixteen participants used hearing aids regularly or sometimes. The majority were born to hearing parents (deaf parents = 6; hearing parents = 15; responded ‘non applicable’=1).

Outside the scanner, both groups completed the Vernon-Warden Reading Test (Kirklees Reading Assessment Schedule) (83). The hearing participants (mean raw score = 34, SD = 4, range = 24-40) scored significantly higher (t (45) = 3.853, p = 3.68 x 10^-4^, d = 1.13) than the deaf participants (mean raw score = 28, SD =7, range = 10-41) on this test. All deaf participants except for one had a reading age older than 13 years of age. Thirteen of the twenty-two participants (59%) had a reading age of 16 or above (the end of compulsory schooling in the UK).

Both groups also completed the Test of Adult Speechreading (TAS) (84) using the minimal pairs, single words, sentences and stories subtests. Deaf participants were on average better than the hearing participants (F (1,45) =31.824, p = 1.00 x 10^-6^, η_p_ ^2^ = 0.41). A significant interaction indicated that this group difference was most pronounced on the Stories subtest (F (3,135) = 2.751, p = 0.045, η_p_ ^2^= 0.06), which placed the greatest demands on speechreading.

There was a positive relationship between speechreading (averaging over the subtests of the TAS) and text reading ability in the deaf participants (r (20) = 0.518, p = 0.007, one tailed test) but not hearing participants (r (23) = −0.240, p = 0.124, one tailed test, SI Appendix, Fig S1).

### Materials

Eight consonant-vowel-consonant words were used as stimuli: beam, beat, boom, boot, real, reef, rule, roof. These words were chosen so that subsets shared initial consonant, vowel and final consonant (see Figure 1C). The phonemes in each position were selected to be maximally visually discriminative when viewed as visual speech and high in frequency and imageability, low in age of acquisition and well matched across psycholinguistic properties (see SI Appendix, SI Methods and Table S2). There was no evidence of a correlation between the Shared Phonemes model (Fig 1C) and theoretical models based on these psycholinguistic properties (all ps >= 0.254).

Speech samples were recorded (audio only) by a male and female speaker. Two different articulations of each word were recorded. The recording of each individual word was filtered to account for the frequency response of the Sensimetric headphones used in the scanner (http://www.sens.com/products/model-s14/) and the overall amplitude was Root Mean Square (RMS) equalised to ensure a similar perceived loudness. The mean duration of the auditory stimuli was 756 ms (min = 558 ms; max = 972 ms; range = 414 ms).

Visual speech exemplars were established by recording audio-visual productions by the same male and female models as the auditory stimuli. The audio was removed. A video sampling rate of 50 fps and an aspect ratio of 1920×1080 was used. Two different articulations of each word were made by each speaker. The head and neck of each speaker was shown within frame against a grey background (see Fig 1A). Each spoken word was produced in isolation with the vocal apparatus returning to a resting state before and after articulation. Videos were sampled to 25 frames per second and a resolution of 960 x 540 with Adobe Premiere for presentation in the scanner. The mean duration of the visual speech articulations was 1243 ms (min = 1040 ms; max = 1360ms; range = 320 ms).

A dynamic text stimulus was generated for each of the eight words. In these videos, each word was revealed sequentially letter-by-letter. The stimuli were animated using Apple Motion. Words were presented in the centre of the screen as white text on a grey background. Four examples of each word were created such that they differed in font type (Nella Sue and Ling Wai TC) and font size (44pt and 48pt). Sequentially delivered stimuli were animated so that individual letters appeared in sequence from left to right, with the strokes comprising each letter appearing in fluid stages (see Fig 1B, right). As later letters appeared, preceding letters faded and disappeared. Elements of two or three letters of each stimulus were visible simultaneously as the item sequentially unfolded. When reading dynamic text, participants are not able see the whole word as the letters appear and disappear sequentially over time, promoting attention to graphemes and their corresponding phonemes. Consistent with this interpretation, sequential delivery engages the dorsal reading pathway, including the posterior superior temporal and inferior parietal cortex, in hearing individuals even for highly familiar words that might ordinarily be read via ventral, lexicosemantic routes (22). The mean duration of the dynamic text was 1195 ms (min = 1000ms, max = 1320 ms; range = 320 ms).

### fMRI Paradigm

#### MRI data acquisition

Data was acquired with a 3-Tesla scanner using a Magnetom TIM Trio systems (Siemens Healthcare, Erlangen, Germany) with a 32 channel headcoil. A 2D epi sequence was used comprising thirty-five 3mm thick slices using a fast-sparse ascending sequence (TR=3950ms, TA=2450ms, FA= 90°, TE=30ms, matrix size= 64×64, in-plane resolution: 3mm × 3mm, interslice gap = 1mm). Fast-sparse acquisition (23) ensured that auditory stimuli could be heard clearly without competing background noise from the scanner (see Fig 1B, right). Stimuli were played during a 1.5s silent gap between volume acquisitions. The same sequence was used for deaf and hearing participants so that data could be combined for the visual speech condition across groups. To ensure there was no forward noise masking from the scanner, stimulus presentation began 50ms after the offset of scanner acquisition.

Four runs of data were acquired each lasting ~12 minutes with 178 brain volumes collected per run (712 volumes per participant). The first three volumes were discarded to account for T1 equilibrium effects. The fourth scan was included as the first modelled scan (e.g. time = 0 seconds), each run began with a ‘rest trial’ (that constituted one of the 10 rest trials) and with five additional scans at the end of each run to allow return to baseline. EPI data collection lasted around 50 minutes. This was followed by a fieldmap, acquired using a double-echo FLASH gradient echo sixty-four slice sequence (TE1=10ms, TE2=12.46ms, in-plane view 192×192 mm, in-plane resolution: 3mm × 3mm, interslice gap = 1mm). At the end of the session a high-resolution T1 weighted structural image was collected using a 3D Modified Driven Equilibrium Fourier Transform (MDEFT) sequence (TR=1393ms, TE=2.48ms, FA= 16°, 176 slices, voxel size = 1 × 1 × 1 mm).

In the scanner, stimuli were presented using the COGENT toolbox running in MATLAB. Auditory stimuli were presented at the same comfortable listening level for all participants.

Visual images were presented with a screen resolution of 1024×768 and a frame rate of 60Hz, using back projection onto a within bore screen at a distance of 62cm from the participants’ eyes.

#### fMRI task - behavioral training and testing

In the scanner, hearing participants were presented with visual only speech and auditory only speech and deaf participants with visual only speech and dynamic text. Prior to entering the scanner, participants were briefly trained to recognise the words presented as visual speech (deaf and hearing) and dynamic text (deaf only) and practiced the within scanner task (described below). After scanning, participants were tested on their ability to identify the words presented as visual speech (both groups) and dynamic text (deaf only) and were highly accurate (see SI Appendix, SI Methods).

#### fMRI task

Participants took part in 4 runs of data collection in total (deaf – visual speech/ dynamic text; hearing – visual speech/ auditory speech). Each run consisted of 170 trials in total. This included 128 stimulus trials, 10 null trials, 16 target and 16 decision trials. Each trial lasted 3950ms (e.g. the time to repetition of each brain volume, inclusive of the silent gap).

A fixation cross was displayed at all times except during stimulus presentation of visual speech and dynamic text, and during the decision trials. For the visual speech and dynamic text trials, the fixation cross was replaced by a video of the dynamic text or lips of the speaker, centred at the location where the fixation cross was previously displayed.

For the hearing group, the eight words were presented as visual speech or auditory speech, produced by a male and a female language model, with two different exemplars (e.g. different recordings) that were repeated twice. For the deaf group, the eight words were presented as visual speech by a male and female language model in two different exemplars and repeated twice and as dynamic text in different fonts and font sizes and repeated twice. For both groups this resulted in 128 trials per run (8 words x 2 stimulus types x 2 language models/fonts x 2 exemplars x 2 repetitions).

During scanning, participants were required to attend to each stimulus and were occasionally asked to identify the word that was presented on the previous trial in a 2-Alternative Forced Choice (2AFC), one-back task memory task (see Fig 1B). These trials constituted 32 trials per run, comprising the target video or sound (16 trials) and a decision trial in which they were instructed to respond (16 trials, 16/128 = 13% of total number of stimulus trials). On these decision trials, participants were cued to respond by a question mark that appeared in the centre of the screen, replacing the fixation cross. To the left and right of that question mark appeared the 2-alternative choices: the target (preceding item) and the foil – both were static written words (see Fig 1B). Participants were required to indicate with a button press whether the correct response was located on the left or right of the screen. The handedness of the button response was counterbalanced across participants. The location of the correct response on the screen, e.g. left or right, was randomised such that it appeared on each side with equal probability across the run. In each run, the target stimuli, constituted all 8 words presented in equal number in each of the two stimulus types (e.g. 8 visual speech and 8 auditory words; 8 visual speech and 8 dynamic text) with an equal and balanced number from the two speakers/font types and exemplars/font size. Over the course of the whole experiment, it was ensured that all 64 unique stimuli were presented once as a target stimulus. The foils were selected randomly from the alternative words. The button press data from one deaf participant was discarded due to error in data collection, as the participant pressed a non-designated button for one of the response options.

Within-run structure was the same for both groups, stimulus trials were randomly permuted in 2 blocks of 64 trials (128 stimulus trials total), each block containing each unique stimulus once. This randomisation was constrained such that the same lexical item regardless of stimulus type could not be presented consecutively to prevent habituation. The 10 null trials, 16 target trials, 16 decision trials were then pseudo-randomly distributed throughout each run, with the constraints that they were relatively evenly distributed (distance between successive null trials: min = 8; max = 26; mean = 17, SD = 3) and that the decision trials had to follow target trials (distance between successive decision/target trials: min = 7; max = 14; mean = 11, SD = 2).

## Statistical analysis

### Univariate Analysis

Univariate analyses were conducted to establish whether the data were consistent with previous univariate studies and the current RSA analyses (see SI Appendix, SI Methods & SI Results).

### Representational similarity analysis (RSA)

Data were analysed with SPM12 (http://www.fil.ion.ucl.ac.uk/spm/) with MATLAB. Functional scans were realigned to the first image and unwarped using field maps. The structural image was co-registered to the mean functional image. The parameters derived from segmentation, using the revised SPM12 segmentation routines, were applied to normalise the functional images to the MNI template at 3mm^3^. The RSA data were not smoothed.

At the first level, data were analyzed using a general linear model with a 350 second high-pass filter and AR(1) autocorrelation model. Events were modelled with a canonical HRF marking the onset of the stimulus and duration in seconds. In the first level model, each word was modelled as a separate regressor (32 regressors: 8 words x 2 stimulus types x 2 forms (e.g. speakers or fonts) with each regressor comprising four trials (2 repetitions x 2 exemplars).

Additional regressors were included modelling the onset of target trials with one regressor for each stimulus type (e.g. onset of the occasional stimulus on which a decision was made), the visual cue to decide and button presses. Target trials were modelled separately so that there were the same number of stimulus trials per regressor. This constituted 36 regressors per run, plus 6 motion parameter regressors, and 4 session means. The rest condition constituted an implicit baseline.

RSA analysis was conducted with the RSA toolbox (85) (https://github.com/rsagroup/rsatoolbox_matlab, version 2.0, accessed March 2017). The beta weights (regression coefficients from the first-level GLM) were used to calculate the cross-validated Mahalanobis (crossnobis) distances (86). First, the beta weights underwent multivariate noise normalisation that down-weights correlated noise across voxels. Then the cross-validated distance was calculated only across separate imaging runs to ensure that the estimated distances between neural patterns are not systematically biased by run-specific noise. This allowed us to test the distances directly against zero (as one would test cross-validated classification accuracy against chance). Therefore, the crossnobis distance provides a measurement on a ratio scale with an interpretable zero value that reflects an absence of distance between items.

For the majority of analyses, a two-step analytic approach was used (SI Appendix, SI Methods for further details). At step (1), searchlight analysis identified regions of interest in which there was significant information (e.g. reliable non-zero crossnobis distances). Then at step (2), in regions shown to contain reliable distances, the distances from the searchlights were averaged and correlated with theoretical models. For one additional analysis in the hearing group in the left STC/MTC (see results section & SI Appendix, Fig S8), we tested directly for fits between the Sublexical model (see below) and representational distances at each searchlight location (rather than adopting the two-step approach explained above).

### RSA Models

We tested three models. The Shared Phonemes model predicted dissimilarity based on the number of shared phonemes, by position, between words (Fig 1C). To test specifically for sublexical encoding of phonology, we fit an additional Sublexical model in which same word-to-same word dissimilarities were not tested (i.e., the diagonal elements of the model were excluded) and dissimilarity based on sublexical structure was predicted on the ‘off diagonal’ of the model only (Fig 1D). This was contrasted with a Lexical model, which predicted that each word is uniquely represented, such that each word is most similar to itself and maximally and equally dissimilar to all other words (Fig 1E).

These models were tested against each set of distances. For hearing participants, we tested in the (1) auditory speech distances, (2) visual speech distances and (3) within-stimulus distances combined across stimulus types (i.e., within auditory speech *and* within visual speech distances) and (4) all across-stimulus (visual speech to auditory speech) distances. For deaf participants, we tested against the (1) dynamic text distances, (2) visual speech distances and (3) within-stimulus distances combined across stimulus types (i.e., within dynamic text *and* within visual speech distances) and (4) all across-stimulus (visual speech to dynamic text) distances. In regions where there was a significant fit in (3), i.e., the combined within-stimulus distances, we further tested (4) for a fit in the *across-stimulus* distances. A fit to both the within and across-stimulus distances provides evidence for shared representational structure across stimuli. We only tested for across stimulus representations in regions showing within-stimulus representation (across both language forms shown to participants) as regions representing structure across stimulus would need to *also* represent within-stimulus information.

The models were Kendall’s Tau-a correlated with representational distances for each participant and the resulting coefficients were converted to a Fisher Z value for group level statistical testing. As negative correlations are not plausible, greater than 0 model fits were assessed with one-tailed, one sample t-tests. Two-tailed paired t-tests were used to assess differences in fit between models.

## Acknowledgments

This research was funded by Wellcome Senior Research Fellowships awarded to MM (100229/Z/12/Z and 220291/Z/20/Z). CP is funded by Wellcome (203147/Z/16/Z, 205103/Z/16/Z and 224562/Z/21/Z). We would like to thank Dr. Kate Rowley for providing comments and helpful discussion on the manuscript. We would also like to thank all participants for their involvement in the study.

