## Supplementary materials for "Reading ability in both deaf and hearing adults is linked to neural representations of abstract phonology derived from visual speech"

**Supporting Information Text**

**SI Methods**

***Prescreening speechreading ability in hearing participants***

Speechreading skills vary greatly between individuals. Potential hearing participants were trained with corrective feedback to recognise the words. They were exposed to the two different exemplars of each word produced by the two different visual speech models (8 words x 2 exemplars x 2 speakers = 32 trials). They were then tested without feedback on the same videos presented twice (8 words x 2 exemplars x 2 speakers x 2 repetitions = 64 trials). They were required to type the word that they saw on a laptop. Only participants scoring >= 90% on this task were invited to take part in the scanning session. Twenty-six of forty-one potential hearing participants (63%) reached this threshold. One participant that achieved this accuracy did not return to take part in the scanning session.

***Exclusion of scanning data based on degree of deafness***

Deaf participants were asked to provide their most recent audiogram. Scanning data were initially collected from 24 deaf participants. Two participants provided audiograms indicating moderate rather than severe or profound deafness (defined as < 70 dBHL in the better ear) after they had already been scanned. The data from these participants was removed from the dataset. Audiograms were not available for five participants, but these participants confirmed verbally that they had received a diagnosis of severe or profound deafness by an audiologist.

***Word selection***

Words, rather than non-words, were used to aid visual speech recognition, as speechreading is aided by lexical constraints (1). Speech is phonetically impoverished when seen without access to sound. For example, the sounds /b/, /p/ and /m/ share the same place of articulation and so are difficult to differentiate by sight alone. These sounds are described as belonging to the same visemic class. We took this reduced distinctiveness into account by choosing words in which each speech segment was visually discriminable by position in each word and so, in combination, each word was uniquely disambiguated from every other word. We used the 12 visemic categories described by (2). The chosen words begin with either /r/ or /b/ (which belong to different visemes), contains the vowel sound /u:/ or /i:/ (which belong to different visemes), or ends with /m/, /t/, /f/ or /l/ (which belong to different visemes). This ensured that models based on visemes or shared auditory phonemes would be the same. For example, ‘boom’ and ‘reef’ differ by 3 phonemes and 3 visemes, ‘boom’ and ‘roof’ differ by 2 phonemes and 2 visemes and ‘boom’ and ‘boot’ differ by 1 phoneme and 1 viseme, and so forth.

***Behavioural performance***

*Pre-scanning tasks*

Prior to scanning, participants were briefly trained to recognize the words presented as visual speech and dynamic text. This was an 8 Alternative Forced Choice Task during which they received corrective feedback (32 trials per input type - visual speech /dynamic text). Following this, they also practiced the scanning task outside the scanner in a truncated version (40 trials) of a single run of data collection.

*Post scanning tasks*

After scanning, participants were tested on their ability to identify the words (deaf/ hearing - visual speech; deaf - dynamic text). Data were not collected for hearing participants for identifying heard speech, given that this would be at ceiling (as it was not degraded). Each video was presented twice (64 trials each for the visual speech and dynamic text stimuli). Participants typed the perceived word into a laptop. Proportion correct data were converted to Rationalised Arcsine Units (RAUs). Both groups were highly accurate on the visual speech comprehension test (see SI Appendix, Fig S2 for the confusion matrices). However, the deaf participants (mean proportion correct = 0.97) performed significantly better ($t(45) = 2.438$, $p = 0.019$, $d = 0.71$) than the hearing participants (mean proportion correct = 0.94). Deaf participants were highly accurate in identifying the dynamic text (mean proportion correct = 0.98).

*Within scanner task*

Accuracy was very high in the within scanner task - hearing group: 97% auditory speech, 92%, visual speech; deaf group – 94% dynamic text, 96% visual speech. The deaf group were more accurate at the visual speech task ($t(44) = 2.861$, $p = 0.006$, $d=0.84$) but the magnitude of difference was numerically small, representing a mean difference of only 4%.

#### ***Univariate analysis methods***

Data were analysed with SPM12. Preprocessing and first level modelling were the same as for the RSA analyses - except that the normalized images entered into the first level model were additionally smoothed with a Gaussian kernel of 8-mm full-width half maximum. Contrast images for [Auditory Speech > Rest], [Visual Speech > Rest] and [Dynamic Text > Rest] and [Hearing: Auditory Speech > Visual Speech] and [Deaf: Dynamic Text > Visual Speech], were taken to the second level to conduct one sample t-tests. Conjunction null analyses were conducted in SPM using the ImCalc function: [(i1 > 0)*(i2>0)] which were conducted on the thresholded statistical maps resulting from the one sample t-tests. All statistical maps were thresholded at an uncorrected peak level threshold of $p < 0.001$, FDR cluster corrected at $q < 0.05$ at the cluster level.

#### ***RSA analytical approach***

*Searchlight RSA (step 1)*

The two-step approach below is described visually in SI Appendix Fig S3. A volumetric searchlight analysis (3) was conducted using a spherical searchlight containing 65 voxels (4). In each searchlight, the crossnobis distance between the neural response to each stimulus and every other stimulus was calculated to generate a Representational Dissimilarity Matrix (RDM) for every voxel and its surrounding neighbours ('a searchlight'). This was estimated at each and every voxel across the brain.

Distances were calculated within each individual stimulus separately (SI Appendix, Fig S3A, lighter or darker grey boxes), within-stimulus types in combination (SI Appendix, Fig S3A, both grey boxes; for example within auditory *and* within visual speech in hearing participants), and across-stimulus type (SI Appendix, Fig S3A, yellow box) at each searchlight location. For the hearing participants, the resulting RDM comprised auditory speech - auditory speech, visual speech - visual speech, and auditory - visual speech distances, which constituted the within and across-stimulus dissimilarities. For the deaf participants, the resulting RDM comprised dynamic text-dynamic text, visual speech-visual speech and dynamic text-visual speech distances, which constituted the within and across-stimulus dissimilarities.

For hearing participants, we ran searchlights to identify: (1) positive auditory speech distances, (2) positive visual speech distances and (3) positive within-stimulus distances (averaging over both auditory and visual speech distances). For deaf participants, we ran searchlights to identify (1) positive dynamic text distances, (2) positive visual speech distances and (3) positive within-stimulus distances (averaging over both dynamic text and visual speech distances). These distances were calculated only between stimuli from the different speakers or fonts (in the case of dynamic text) to focus on abstract representations, excluding similarities driven by low-level perceptual properties, as cross-validation of the distances ensures that positive distances demonstrate representation of the respective stimuli. In each group, we averaged the distances at each searchlight location, returning the averaged distance value to the voxel at the centre of each sphere. Distances were averaged at each searchlight location for each separate stimulus (SI Appendix, Fig 3B, lighter or darker grey boxes) or combined within-stimulus type (SI Appendix, Fig 3B, both grey boxes) depending on the analysis.

These distances were then submitted to a whole brain one sample t-test in which each voxel constituted the average distance at a searchlight location. Each participants' whole brain searchlight map was inclusively masked with a >20% probability grey matter mask, using the canonical MNI brain packaged with SPM12. The masked images from each participant were submitted to SPM12 for one sample t-tests testing for reliable non-zero distances (e.g. > 0) and a two-sample t-test to assess differences between groups in the distances for visual speech. All statistical maps are presented at an uncorrected peak level threshold of $p < 0.001$, FDR corrected at $q < 0.05$ at the cluster, unless stated. Resulting clusters from these analyses were used to identify regions of interest for subsequent analysis.

*Regions of Interest (ROI) Analyses (step 2)*

ROI analyses are advised when testing special populations in which sample sizes are necessarily restricted (5). It is important that the RSA models are evaluated within regions of interest that were defined in a manner that is statistically unbiased (6). Note that when non-zero stimulus distances are used to identify ROIs, the ROI definition and evaluation of the model are independent as the mean distance is implicitly subtracted out in the correlation between the model and the distances.

The searchlight procedure identified regions of interest (clusters) containing significant information (i.e., regions with reliable non-zero distances). In these regions, the response of each region was summarised by averaging the distances at all the searchlight locations that comprised each cluster (SI Appendix, Fig S3C). This was conducted on each set of individual stimulus distances, the combined within-stimulus distances or across-stimulus type depending on the analysis. The resulting distances were correlated with theoretical models (Fig 1C-E). The non-parametric Kendall's Tau-a rank correlation was used in these analyses in preference to Pearson or Spearman correlations as the models contained tied ranks (7). The resulting correlation coefficient was converted to a Pearson's r value, then to a Fisher-transformed Z value, to permit parametric statistical analysis (8).

Multidimensional Scaling (MDS) was conducted to visualise the similarity structure of the RDMs. Given that univariate activity was on average greater for auditory speech relative to visual speech in hearing participants and greater for visual speech relative to dynamic text for deaf participants in the STC/MTC (see Fig S6 A & D), the first dimension, which reflected this univariate difference in overall signal magnitude was removed, from the MDS visualisations.

### SI Results

#### *Univariate control analyses*

We conducted a set of control analyses with the purpose of confirming that our fMRI data were consistent with previous studies and were broadly consistent with the RSA analyses (SI Appendix, Fig S4 A-D). As expected, these analyses showed significant overlap in activity relative to rest in the bilateral STC and/or MTC across language conditions and groups (SI Appendix, Fig S5 A-C). Whilst activity was co-located in the STC and/or MTC for different language stimuli, the magnitude of activity was greater for auditory speech than visual speech in hearing participants, and for visual speech compared to dynamic text in deaf participants, in these regions (SI Appendix, Fig S6 A-D). Activation tables for these analyses are available via the following OSF link here: https://osf.io/cba8z/overview.

#### *RSA analyses within each individual stimulus condition*

Prior to conducting the main analyses aimed at isolating across-stimulus type encoding of phonology, we tested for neural representations of phonology that were abstracted from lower-level sensory features within each individual language stimulus (i.e., for auditory speech, visual speech and dynamic text separately). To do this, we estimated activity patterns elicited by each trial and calculated the distance between patterns within searchlight locations across the whole brain. We identified clusters with non-zero distances in the across speaker or across font distances for visual speech, auditory speech and dynamic text separately (SI Appendix, Fig S7A). In clusters with non-zero distances, we then assessed the fit to the Shared Phonemes model (Fig 1C). If there was a fit to the Shared Phonemes model, we additionally tested for a fit to the Sublexical (Fig 1D) and Lexical models (Fig 1E) to isolate sublexical from lexical representation within that region. Activation tables for the resulting searchlight maps are available via the OSF link above.

In the hearing group perceiving auditory speech we identified 3 clusters with reliable non-zero distances (SI Appendix, Fig S7B). This included clusters in the left lateral STC/MTC [peak at -54 -10 5], right STC/MTC [57 -31 5] and a more medial cluster in the left posterior STC [-42 -31 -1]. In all three clusters, there was a significant fit to the Shared Phonemes model and the Sublexical model (SI Appendix, Table S3).

In the deaf group perceiving dynamic text searchlight analyses identified 6 clusters with reliable non-zero distances (SI Appendix, Fig S7C). These were focused mainly in the bilateral temporal and occipital cortices. There was a fit for the Lexical model in the left lateral occipital complex (SI Appendix, Table S3), but not to either the Lexical or Sublexical model in the left STC/MTC cluster. However, when we reduced the peak level threshold to $p < 0.005$ uncorrected, the cluster in the left STC/MTC was much larger (peak at [-57 -25 4], 74 voxel extent, $Z = 4.19$) and there was a significant fit to the Sublexical model ($t(21) = 2.76$, $p = 0.006$, $dz = 0.59$) at an uncorrected level $p < 0.05$.

For the visual speech condition (both groups), there were no significant differences between groups in the searchlight maps when these were directly tested and so we collapsed the data across the deaf and hearing group. Searchlight analyses identified 11 clusters with positive visual speech distances. These were focused mainly in left middle temporal and temporoparietal cortex, bilateral inferior frontal cortex, left middle frontal gyrus and bilateral occipitotemporal cortex (SI Appendix, Fig S7D). Only two clusters were a significant fit to the Shared Phonemes model (SI Appendix, Table S3). In the left MTC cluster [peak at -54 -31 -10], there was a significant fit to the Sublexical model. Whilst in a more posterior and superior cluster in the left temporo-parietal cortex [peak at -57 -52 11] there was a fit to the Lexical model only. Note that there were no significant differences between groups in the model fits (all $ps \geq 0.249$).

Taken together, these within-stimulus analyses provided evidence of sublexical representations of phonology in the STC/MTC for each stimulus type individually, albeit this evidence was less robust for representations of phonology in dynamic text for deaf participants.

### Figures

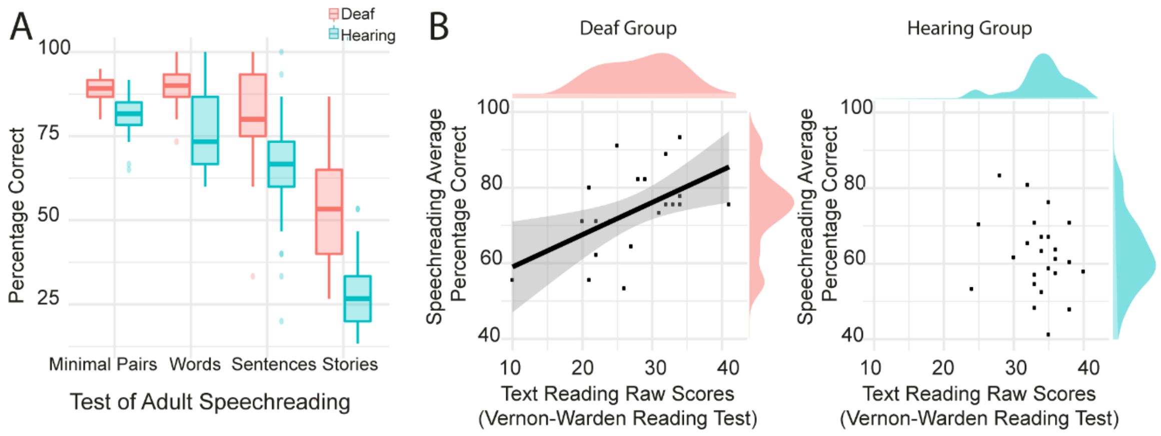

**Fig. S1.** Speechreading in deaf and hearing participants and the relationship to text reading. (A) Participants completed the Test of Adult Speechreading (TAS) outside the scanner. This showed that deaf participants were better speechreaders than hearing participants and this difference was most pronounced in the Stories subtest. (B) Deaf participants who were better speechreaders were better text readers (left panel). There was no evidence of a relationship between speechreading and text reading in the hearing participants (right panel). Reading scores were on average lower in the deaf than hearing group.

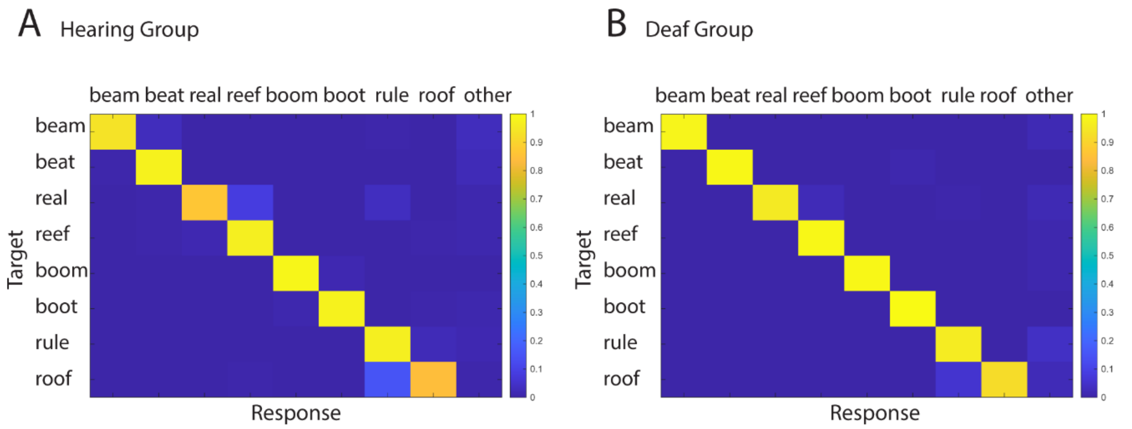

**Fig. S2.** Confusion matrices for the post-scanning visual speech identification task based on typed response. (A) Confusion matrices for the Hearing Group and (B) Confusion matrices for the Deaf Group. If the typed response was different to the possible targets, this was marked as other.

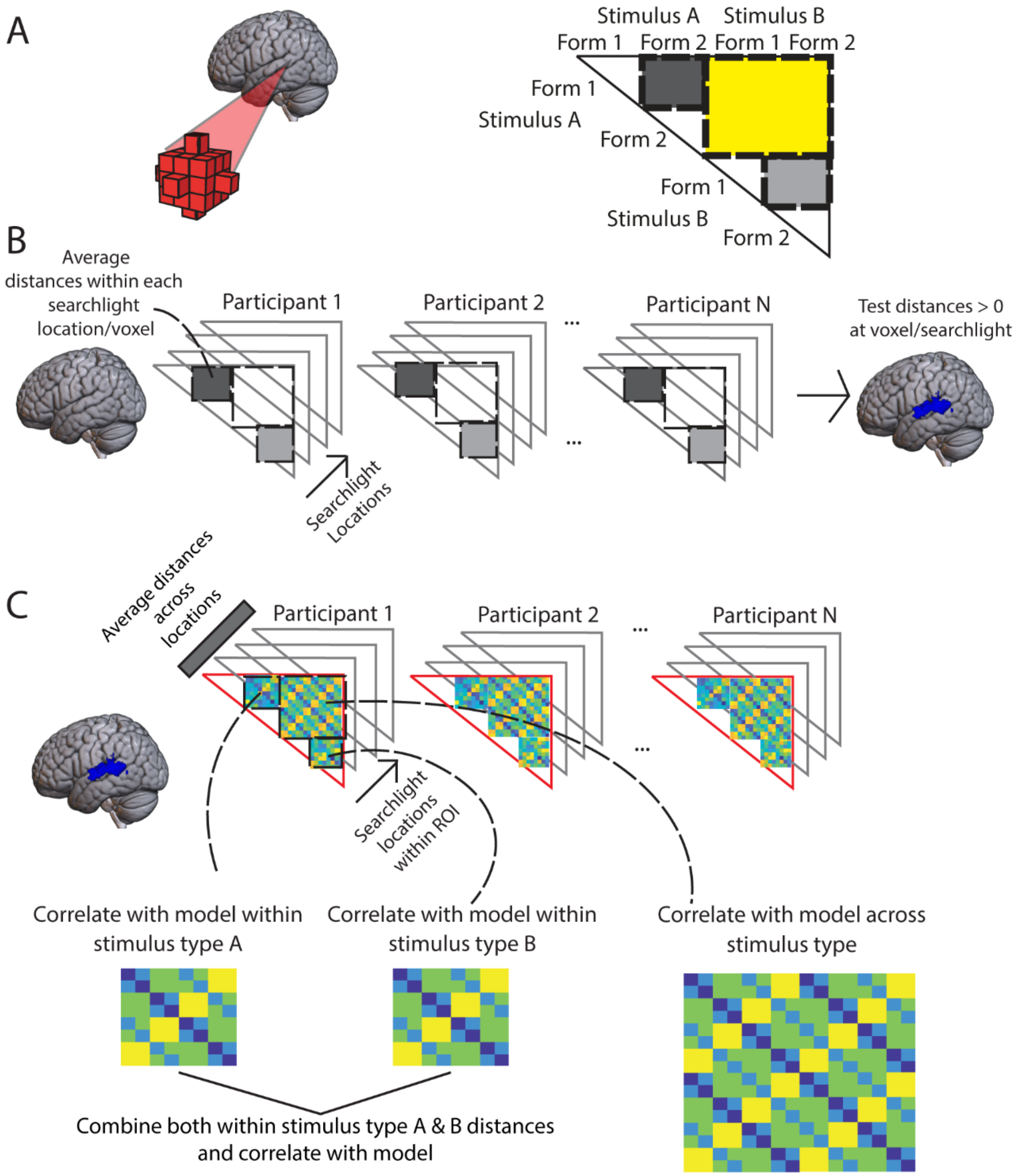

**Fig. S3.** Description of the Analytic approach for the Representational Similarity Analysis. A) Representational distances were calculated within searchlights covering the whole brain. Distances were calculated across speaker/fonts within each individual stimulus separately (lighter or darker grey boxes), within-stimulus types in combination (both grey boxes) and across-stimulus type (yellow box) at each searchlight location. B) To identify ROIs for subsequent analyses, representational distances were averaged at each individual searchlight location within each individual stimulus separately (lighter or darker grey boxes) or within-stimulus types in combination (both grey boxes) depending on the analysis. These searchlight maps were then submitted to a whole brain analysis to identify regions of interest with > 0 distances. As these distances were cross-validated, positive distances demonstrated representation of the respective stimuli. C) Searchlight distances were extracted from regions of interest/clusters with > 0 distances, the appropriate set of distances (i.e. within individual stimulus, within both stimulus types or across-stimulus type) were then averaged across searchlights/locations within each region/cluster to summarise the response in each participant, and the averaged distances were then correlated with theoretical models (i.e. the Shared Phonemes, lexical and sublexical models). The correlation coefficients for each participant were converted to Fisher Z values for group level statistical analyses.

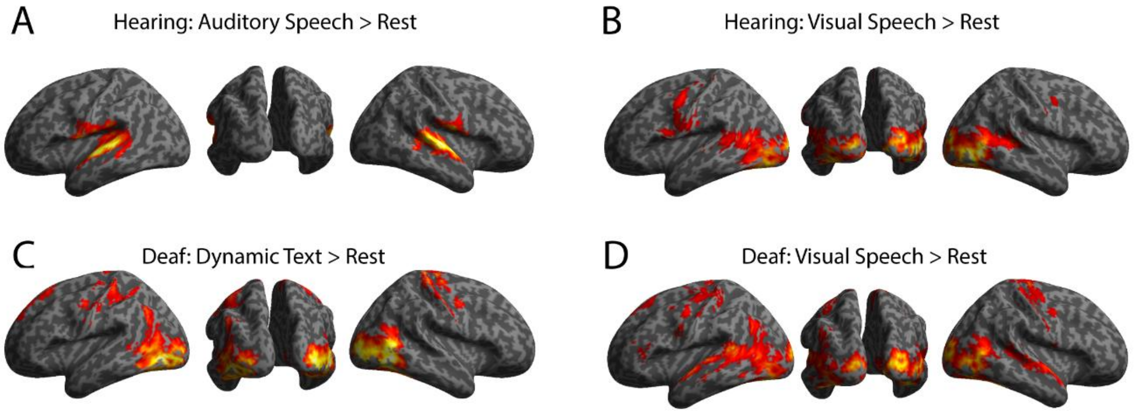

**Fig. S4.** Univariate analyses of each condition relative to rest rendered on the inflated MNI brain and thresholded at $p < 0.001$ peak uncorrected, $q < 0.05$ FDR corrected at the cluster level. (A) In the hearing group, [Auditory Speech > Rest] and (B) [Visual Speech > Rest]. (C) In the deaf group, [Dynamic Text > Rest] and (D) [Visual Speech > Rest].

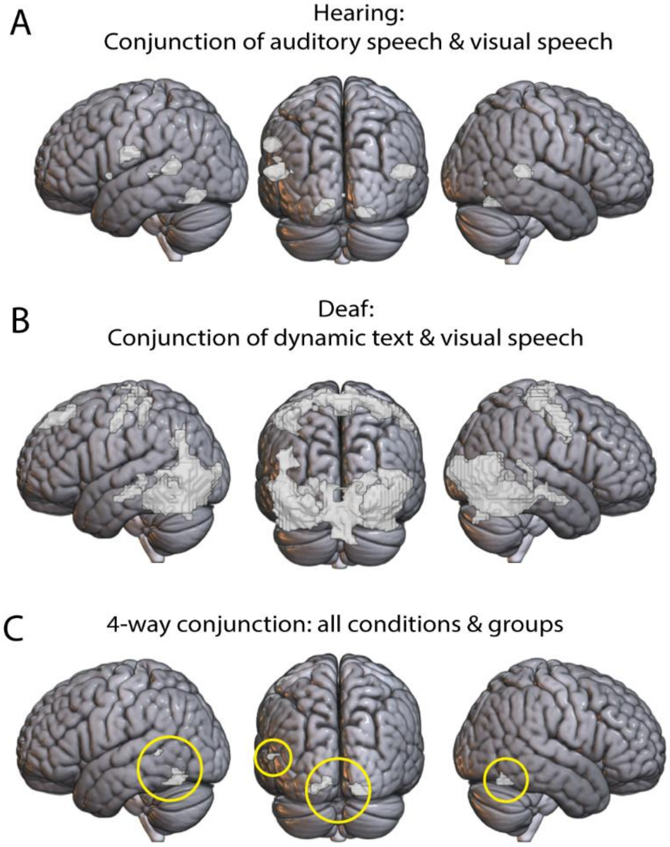

**Fig S5**. Univariate conjunction null analyses rendered on the single subject MNI brain and thresholded at $p < 0.001$ peak uncorrected, $q < 0.05$ FDR cluster corrected. (A) Conjunction of [Auditory Speech > Rest] and [Visual Speech > Rest] in hearing participants. (B) Conjunction of [Dynamic Text > Rest] and [Visual Speech > Rest] in deaf participants. (C) Conjunction between all conditions and groups, highlighted in yellow circles.

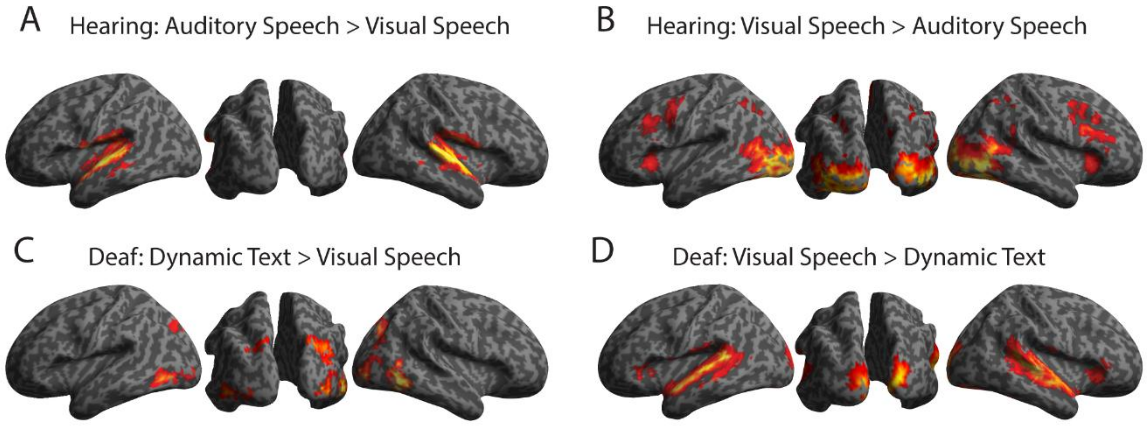

**Fig S6.** Univariate analyses of the between condition comparisons for each group rendered on the inflated MNI brain and thresholded at $p < 0.001$ peak uncorrected, $q < 0.05$ FDR corrected at the cluster. In the hearing group, (A) [Auditory Speech > Visual Speech] and (B) [Visual Speech > Auditory Speech]. In the deaf group, (C) [Dynamic Text > Visual Speech] and (D) [Visual Speech > Dynamic Text].

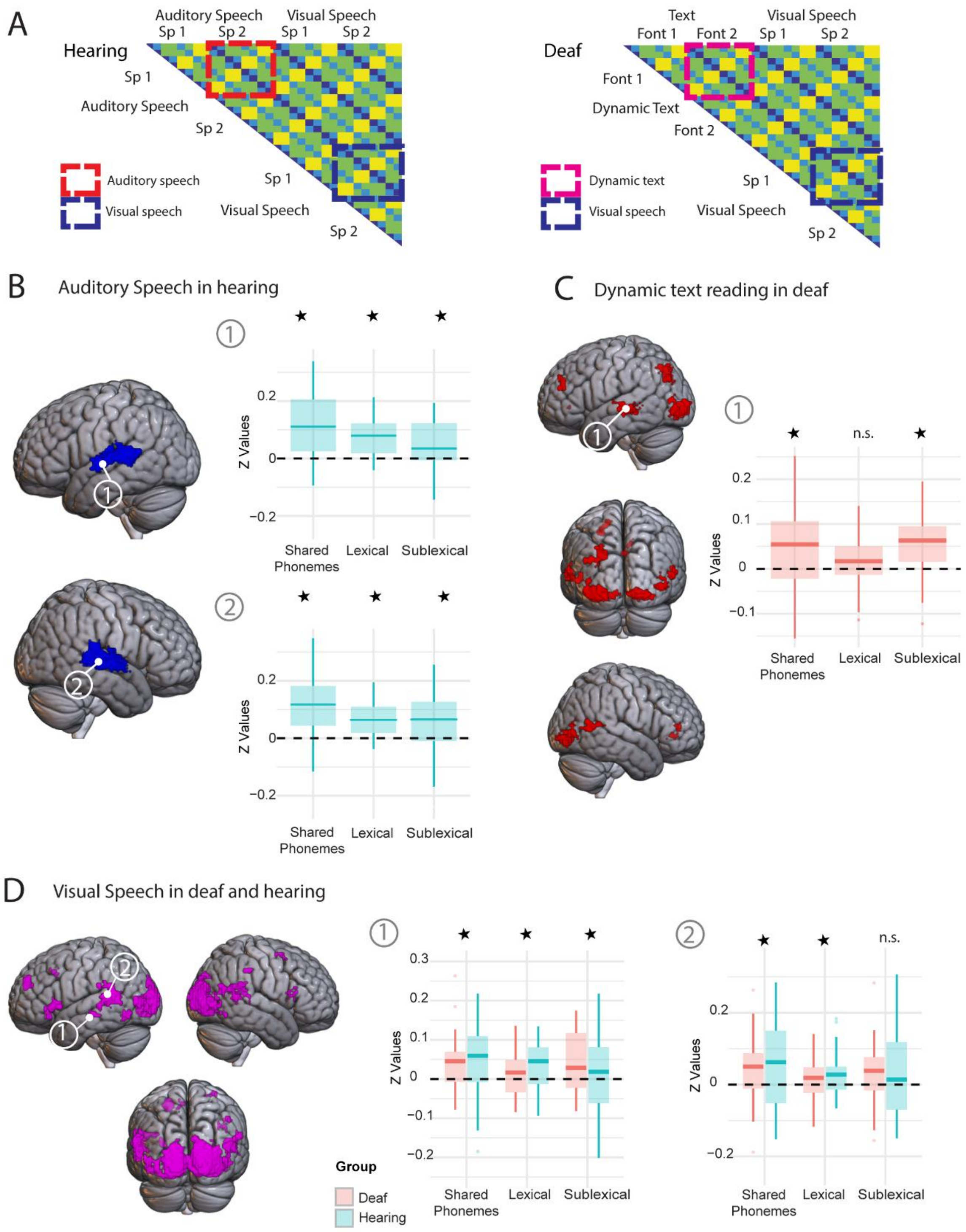

**Fig S7.** Searchlight maps showing clusters with reliable non-zero distances for each individual stimulus, rendered on the MNI brain and thresholded at $p < 0.001$ peak uncorrected, $q < 0.05$ FDR cluster corrected, except in C (see below). (A) Illustration of extraction of stimulus condition distances for hearing and deaf participants. (B) Searchlight map and boxplots of model fits in the STC/MTC clusters for auditory speech in hearing participants. Note for brevity we only show the left lateral STC/MTC and right STC/MTC plots. (C) Searchlight map and boxplots for model fits in the dynamic text distances in the left STC/MTC deaf participants at a reduced threshold of $p < 0.005$ peak uncorrected, $q < 0.05$ FDR cluster corrected. (D) Searchlight map and boxplots for model fits in the visual speech distances collapsing across groups in the two clusters in the left STC/MTC. *****Indicate significance corrected for multiple comparisons across number of clusters/tests, except in S7C in which this indicates uncorrected significance at $p < 0.05$.

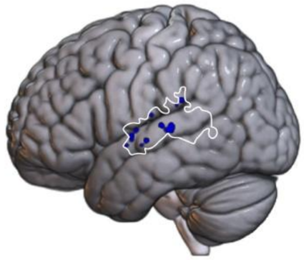

**Fig S8.** Searchlight conducted within the left STC/MTC directly searching for fits between the Sublexical model and the across-stimulus distances in hearing participants. Clusters of voxels significant at $p < 0.005$ peak level uncorrected are shown in blue. Boundary of search volume is shown in white.

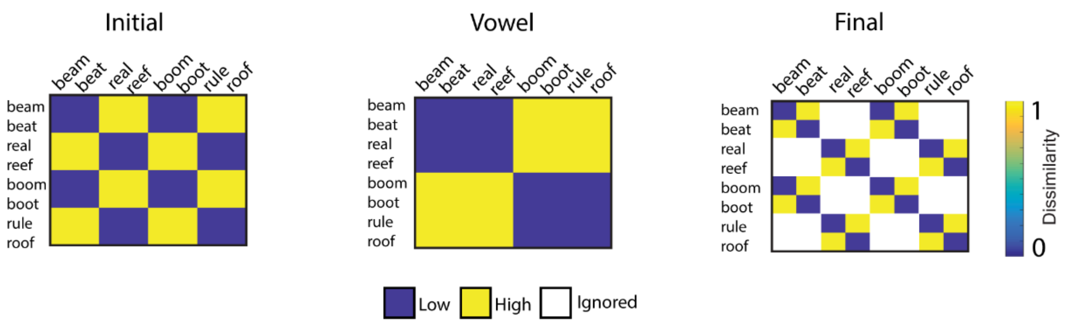

**Fig S9.** Models used to test for differential strength in encoding of abstract phonemes in different segment positions. From left to right, models test for shared initial consonant, shared vowel and shared final consonant.

**Table S1.** Background details of deaf participants. Language and hearing background including dBHL level for best ear is shown, where available. Audiograms were unavailable for 5 participants, but all participants reported that they were severely or profoundly deaf.

| Age of onset of deafness | Aetiology of deafness | Best-Ear dBHL confirmed by audiogram | Self-reported level of deafness | Self-rated BSL proficiency (1-7) | Age (in yrs) started learning BSL | Do you wear hearing aids (Yes/No/Sometimes/Not anymore)? | Preferred language (British Sign Language, BSL; Sign Supported English, SSE; English) | Hearing status of parents (both) |
|---|---|---|---|---|---|---|---|---|
| Born deaf | Genetic | 100 (L) | Profound | 7 | <3 | Yes | BSL | Deaf |
| Born deaf | Genetic | 91 (L) | Severe-Profound | 5 | Don’t know | Yes | SSE | Hearing |
| Born deaf | Genetic | 108 (R) | Profound | 6 | >5 | Yes | SSE | Hearing |
| Born deaf | Unknown | 82 (R) | Severe | 5 | >5 | Yes | English | N/A |
| Born deaf | Genetic | 112 (R) | Profound | 7 | >5 | Yes | BSL | Hearing |
| Born deaf | German measles | 108 (L) | Profound | 4 | >5 | Yes | SSE | Hearing |
| Born deaf | Unknown | 104 (R) | Profound | 6 | >5 | Sometimes | BSL | Hearing |
| Born deaf | Genetic | 96 (L) | Profound | 7 | Birth | Yes | BSL | Deaf |
| Born deaf | Genetic | Confirmed as severe-to-profound by audiologist letter | Severe-Profound | 7 | Birth | Not anymore | BSL | Deaf |
| Born deaf | Unknown | 82 (R) | Severe | 5 | <5 | Yes | English | Hearing |
| Born deaf | German measles | 112 (R) | Profound | 7 | <5 | Sometimes | BSL | Hearing |
| Under 3yrs | Unknown | 95 (L) | Severe-Profound | 7 | >5 | No | BSL | Hearing |
| Under 3yrs | Unknown | Unavailable | Profound | 4 | >5 | Not anymore | SSE | Hearing |
| Born deaf | Genetic | 70 (L) | Severe | 7 | Birth | Yes | BSL | Deaf |
| Born deaf | Genetic | 90 (R) | Severe | 5 | Birth | Not anymore | SSE | Deaf |
| Born deaf | Unknown | Unavailable | Severe-Profound | 7 | >5 | Yes | BSL | Hearing |
| Born deaf | Unknown | 107 (L) | Profound | 6 | >5 | Yes | BSL | Hearing |
| Born deaf | Genetic | 94 (L) | Severe-Profound | 7 | Birth | Sometimes | BSL | Deaf |
| Born deaf | Unknown | Unavailable | Profound | 5 | >5 | Yes | SSE | Hearing |
| Born deaf | German measles | 85 (R) | Severe | 2 | >5 | Yes | English | Hearing |
| Born deaf | Unknown | Unavailable | Profound | 7 | <5 | No | BSL | Hearing |
| Born deaf | Unknown | 115 (R) | Profound | 5 | <5 | No | English | Hearing |

**Table S2:** Psycholinguistic properties of the words. Log-spoken word frequency (CELEX), phonological neighbourhood and imageability (Bristol/MRC) were extracted from the N-Watch program (9) and Age of acquisition from Kuperman et al. (10). Visual speech transcription uses the 12 visemes/phonemic equivalence classes, applying a number for each class based on the order in which they are presented in Table 1 in Auer & Bernstein (2)

| **Orthography** | **IPA transcription** | **Visemes/Phonemic Equivalence Class** | **Frequency (log10)** | **Phonological neighbourhood** | **Imageability** | **Age of acquisition** |
|---|---|---|---|---|---|---|
| boom | bu:m | [6][1][6] | 1.15 | 21 | 449 | 5.56 |
| boot | bu:t | [6][1][9] | 0.61 | 26 | 604 | 3.89 |
| rule | ru:l | [10][1][8] | 1.83 | 20 | 415 | 4.72 |
| roof | ru:f | [10][1][7] | 1.21 | 16 | 604 | 5.00 |
| beam | bi:m | [6][3][6] | 1.27 | 17 | 539 | 8.78 |
| beat | bi:t | [6][3][9] | 1.43 | 24 | 406 | 6.15 |
| real | ri:l | [10][3][8] | 2.47 | 7 | 313 | 4.95 |
| reef | ri:f | [10][3][7] | 0.00 | 15 | 485 | 9.72 |

**Table S3:** Model fits for all clusters with reliable distances for auditory speech (in hearing group), dynamic text (in deaf group) and visual speech (deaf and hearing combined). Tests for fits to the Sublexical and Lexical model were only conducted if there was a significant fit to the Shared Phonemes model. Bonferroni correction was conducted for the number of clusters in which a model was tested. ***Indicates statistically significant p-values corrected for multiple comparisons.**

| **Cluster** | **Shared Phonemes model** | | | **Sublexical Phonemes \| Lexical** | | | **Lexical Only** | | |
|---|---|---|---|---|---|---|---|---|---|
| | t-value (df) | p | $d_z$ | t-value (df) | p | $d_z$ | t-value (df) | p | $d_z$ |
| **Auditory Speech** | | | | | | | | | |
| (1) Left lateral STC/MTC [-54 -10 5] | **5.23 (24)** | **1.16 x $10^{-5}$*** | **1.05** | **2.46 (24)** | **0.011*** | **0.49** | **5.69 (24)** | **3.71 x$10^{-6}$*** | **1.14** |
| (2) Left medial posterior STG [-42 -31 -1] | **2.30 (24)** | **0.015*** | **0.46** | **2.39 (24)** | **0.013*** | **0.48** | 1.55 (24) | 0.068 | 0.31 |
| (3) Right STC/MTC [57 -31 5] | **4.91 (24)** | **2.58 x $10^{-5}$*** | **0.98** | **3.13 (24)** | **0.002*** | **0.63** | **5.28 (24)** | **1.03 x $10^{-5}$*** | **1.06** |
| **Dynamic Text** | | | | | | | | | |
| (1) Left Lateral occipital complex [-42 -82 -4] | **3.04 (21)** | **0.003*** | **0.65** | 1.72 (21) | 0.050 | 0.37 | **2.59 (21)** | **0.009*** | **0.55** |
| (2) Right V1-V3 [15 -79 -4] | 1.34 (21) | 0.098 | 0.29 | | | | | | |
| (3) Left middle and superior occipital gyrus [-24 -73 38] | -0.52 (21) | 0.304 | 0.11 | | | | | | |
| (4) Right middle occipital gyrus [48 -76 2] | 0.28 (21) | 0.390 | 0.06 | | | | | | |
| (5) Left MTC [-57 -25 -4] | 1.44 (21) | 0.082 | 0.31 | | | | | | |
| (6) Right MTC [57 -61 8] | 0.53 (21) | 0.300 | 0.11 | | | | | | |
| **Visual Speech** | | | | | | | | | |
| 1) Left MTC [-54 -31 -10] | **3.42 (46)** | **6.62 x $10^{-4}$ *** | **0.50** | **2.29 (46)** | **0.013 *** | **0.33** | **2.46 (46)** | **0.009 *** | **0.36** |
| 2) Left temporo-parietal cortex [-57 -52 11] | **2.97 (46)** | **0.002 *** | **0.43** | 1.95 (46) | 0.029 | 0.28 | **2.62 (46)** | **0.006 *** | **0.38** |
| 3) Bilateral V1-V3 [18 -97 14] | 1.00 (46) | 0.161 | 0.15 | | | | | | |
| 4) Left inferior frontal cortex [-51 20 2] | 1.04 (46) | 0.152 | 0.15 | | | | | | |
| 5) Left middle frontal gyrus [-21 50 26] | 0.15 (46) | 0.440 | 0.02 | | | | | | |
| 6) Left inferior temporal cortex [-45 -64 -7] | 0.68 (46) | 0.249 | 0.10 | | | | | | |
| 7) Right inferior frontal cortex [42 14 14] | 1.18 (46) | 0.122 | 0.17 | | | | | | |
| 8) Left insula [-30 23 -10] | -0.10 (46) | 0.459 | 0.01 | | | | | | |
| 9) Left superior medial and middle cingulate gyrus [-6 20 41] | -0.13 (46) | 0.448 | 0.02 | | | | | | |

| | | | |
|---|---|---|---|
| 10) Right middle and superior frontal [33 5 59] | 1.75 (46) | 0.043 | 0.26 |
| 11) Right supramarginal gyrus [45 -31 41] | 0.46 (46) | 0.324 | 0.07 |

**Table S4:** Clusters with positive searchlight distances within-stimulus type in hearing participants (i.e., combining visual and auditory speech distances) and for deaf participants (i.e., combining visual speech and dynamic text distances). Thresholded at p < 0.001 peak level, q < 0.05 FDR cluster corrected. Cluster forming extent threshold = 10 voxels (hearing group) and 12 voxels (deaf group).

| **Cluster** | **x** | **Y** | **z** | **Extent** | **Z Value** |
|---|---|---|---|---|---|
| **Hearing group** | | | | | |
| **[1] Left STC/MTC** | | | | | |
| Left STG | -54 | -10 | 5 | 243 | 4.99 |
| Left MTG | -60 | -40 | 8 | | 4.88 |
| Left STG | -54 | -19 | 11 | | 4.75 |
| **[2] Right STC/MTC** | | | | | |
| Right STG | 57 | -31 | 5 | 355 | 5.78 |
| Right STG | 57 | -10 | 2 | | 5.16 |
| Right STG | 63 | -22 | 8 | | 5.01 |
| **[3] Right V1-V3** | | | | | |
| Right cuneus | 21 | -94 | 14 | 129 | 5.11 |
| Right Calcarine Gyrus | 9 | -88 | -1 | | 4.65 |
| Right superior occipital gyrus | 18 | -94 | 23 | | 4.35 |
| **[4] Left inferior frontal gyrus** | -51 | 17 | 2 | 10 | 4.28 |
| **[5] Right posterior middle temporal** | | | | | |
| Right MTG | 45 | -49 | 8 | 11 | 4.21 |
| Right MTG | 48 | -49 | 17 | | 4.09 |
| **[6] Left V1-V3** | | | | | |
| Left superior occipital gyrus | -9 | -100 | 5 | 32 | 3.84 |
| Left calcarine gyrus | -6 | -91 | 11 | | 3.75 |
| Left calcarine gyrus | -6 | -94 | -4 | | 3.50 |
| **Deaf group** | | | | | |
| **[1] Left STC/MTC** | | | | | |
| Left STG | -51 | -43 | 14 | 82 | 4.47 |
| Left MTG | -60 | -25 | -1 | | 4.19 |
| Left MTG | -57 | -34 | 2 | | 3.98 |
| **[2] Bilateral V1-V3** | | | | | |
| Right lingual gyrus | 18 | -85 | -4 | 970 | 5.87 |
| Right cuneus | 18 | -97 | 14 | | 5.36 |
| Left calcarine gyrus | -9 | -97 | 2 | | 5.28 |
| **[3] Right STC** | 57 | -40 | 23 | 24 | 4.57 |
| **[4] Right MTC** | | | | | |
| Right MTG | 57 | -52 | 11 | 37 | 4.34 |
| Right MTG | 54 | -58 | 2 | | 3.60 |
| Right STG | 48 | -46 | 20 | | 3.29 |
| **[5] Right inferior frontal gyrus** | | | | | |

| | | | | | |
|---|---|---|---|---|---|
| Right inferior frontal gyrus | 54 | 26 | 2 | 12 | 4.05 |
| Right inferior frontal gyrus | 45 | 26 | 2 | | 3.26 |
| **[6] Left precuneus** | -3 | -73 | 50 | 13 | 3.96 |
| **[7] Left inferior frontal gyrus** | -48 | 20 | -4 | 14 | 3.70 |
| **[8] Right middle occipital gyrus** | | | | | |
| Right middle occipital gyrus | 45 | -76 | 2 | 14 | 3.52 |
| Right middle occipital gyrus | 39 | -85 | 2 | | 3.50 |

**Table S5:** Model fits for within (visual and auditory speech – hearing; visual speech and dynamic text – deaf) and across stimulus distances. Tests for fits to the Shared Phonemes model were only conducted in the across-stimulus distances, if there was a fit to the Shared Phonemes model in the within-stimulus distances. The Sublexical and Lexical models were only tested in the across-stimulus distances if there was a fit to the Shared Phonemes model in the across stimulus distances. Bonferroni correction was conducted for the number of clusters in which a model was tested.
***Indicates significant fit accounting for multiple comparisons.**

| | **Within-Stimulus** | | | **Across-Stimulus** | | | | | | | | |
|---|---|---|---|---|---|---|---|---|---|---|---|---|
| **Cluster** | **Shared Phonemes model** | | | **Shared Phonemes model** | | | **Sublexical Shared Phonemes \| Lexical** | | | **Lexical only** | | |
| | **t-value (df)** | **p** | **$d_z$** | **t-value (df)** | **p** | **$d_z$** | **t-value (df)** | **p** | **$d_z$** | **t-value (df)** | **p** | **$d_z$** |
| **Hearing participants** | | | | | | | | | | | | |
| [1] Left STC/MTC [-54 -10 5] | **4.79 (24)** | **3.54 x $10^{-5}$ *** | **0.96** | **2.46 (24)** | **0.011 *** | **0.49** | 0.029 (24) | 0.489 | 0.01 | **3.52 (24)** | **8.75 x $10^{-4}$ *** | **0.70** |
| [2] Right STC/MTC [57 -31 5] | **4.61 (24)** | **5.55 x $10^{-5}$ *** | **0.92** | **3.57 (24)** | **7.70 x $10^{-4}$ *** | **0.71** | **2.71 (24)** | **0.006 *** | **0.54** | **2.55 (24)** | **0.009 *** | **0.51** |
| [3] Right V1-V3 [21 -94 14] | 0.70 (24) | 0.244 | 0.14 | | | | | | | | | |
| [4] Left inferior frontal gyrus [-51 17 2] | 0.11 (24) | 0.459 | 0.02 | | | | | | | | | |
| [5] Right posterior MTG [45 -49 8] | 0.71 (24) | 0.244 | 0.14 | | | | | | | | | |
| [6] Left V1-V3 [-9 -100 5] | 0.56 (24) | 0.291 | 0.11 | | | | | | | | | |
| **Deaf participants** | | | | | | | | | | | | |
| [1] Left STC/MTC [-51 -43 14] | **3.95 (21)** | **3.68 x $10^{-4}$ *** | **0.84** | **3.66 (21)** | **7.30 x $10^{-4}$ *** | **0.78** | **3.38 (21)** | **0.001 *** | **0.72** | **1.88 (21)** | **0.037 *** | **0.40** |
| [2] Bilateral V1-V3 [18 -85 -4] | 1.70 (21) | 0.052 | 0.36 | | | | | | | | | |
| [3] Right STC [57 -40 23] | 0.14 (21) | 0.444 | 0.03 | | | | | | | | | |
| [4] Right MTC [57 -52 11] | -0.26 (21) | 0.398 | 0.06 | | | | | | | | | |

| | | | |
|---|---|---|---|
| [5] Right inferior frontal [54 26 2] | 2.27 (21) | 0.017 | 0.43 |
| [6] Left precuneus [-3 -73 50] | 0.82 (21) | 0.468 | 0.17 |
| [7] Left inferior frontal gyrus [-48 20 -4] | 0.70 (21) | 0.245 | 0.15 |
| [8] Right middle occipital gyrus [45 -76 2] | 2.02 (21) | 0.028 | 0.43 |
